# A toxin-mediated policing system in *Bacillus* improves population fitness via penalizing non-cooperating phenotypic cheaters

**DOI:** 10.1101/2022.05.14.491907

**Authors:** Rong Huang, Jiahui Shao, Zhihui Xu, Yuqi Chen, Yunpeng Liu, Dandan Wang, Haichao Feng, Weibing Xun, Qirong Shen, Nan Zhang, Ruifu Zhang

## Abstract

Microbial cooperation is vulnerable to exploitation by social cheaters. Although the strategies for controlling genotypic cheaters have been well investigated, the mechanism and significance of preventing phenotypic cheating remain largely unknown. Here, we revealed the molecular mechanism and ecological significance of a policing system for punishing phenotypic cheaters in the community of a plant beneficial strain *Bacillus velezensis* SQR9. Coordinated activation of extracellular matrix (ECM) production and autotoxin bacillunoic acids (BAs) biosynthesis/self-immunity, punished public goods-nonproducing cheaters in strain SQR9’s community. Spo0A was identified to be the co-regulator for triggering both ECM production and BAs synthesis/immunity, which activates acetyl-CoA carboxylase (ACC) to produce malonyl-CoA, an essential precursor for BAs biosynthesis, thereby stimulating BAs production and self-immunity. Elimination of phenotypic cheaters by this policing system, significantly enhanced population fitness under different stress conditions and in plant rhizosphere. This study provides insights into our understanding of maintenance and evolution of microbial cooperation.

## Introduction

Cooperative interactions are not restricted to complex, higher organism, but also prevalent among microbial communities in many contexts^1,2^. Production of costly public goods that can be used by any cells in a population, is a common cooperative behavior consistently found in diverse microorganisms^3^. Typical public goods include extracellular enzymes for substances digesting^4^, siderophore for iron-scavenging^5^, matrix components for biofilm formation^6,7^, biosurfactants for cooperative swarming^8,9^, and so on. Intriguingly, the considerable cost for producing public goods usually raises cheating individuals in the evolution of cooperation, who contributes no or just a little of their share of the common good^3,4,10^. Therefore, cheaters will have a fitness advantage over fully participating cooperators, and their frequency will increase rapidly, eventually leading to the collapse of cooperative behavior^11^. This “tragedy of the commons” is predicated by natural selection and game theory^12,13^, and has been widely illustrated in various cooperation systems^14,15^.

Despite the exploitation of public goods by cheating individuals, cooperation principally survives cheating during the evolutionary history^16^. Several mechanisms have been proposed to play significant roles in maintaining cooperation by preventing cheater invasion^3,16,17^, mainly including kin selection/discrimination that selectively direct cooperation to genetic relatives^18,19^, facultative cooperation regulated by quorum-sensing (QS) system^20^ or nutrient fitness cost^21^, coupling production of public and private goods^22^, punishment of cheating individuals by cooperator-produced antibiotics^10,23^, partial privatization of public goods under certain conditions^24,25^, and spatial structuring to surround the producers more likely by other cooperators^26^. In general, the emergency of multiple sanction strategies is a consequence of natural selection, which suppress social cheaters and promote public goods production, thereby maintaining microbial community stability and improving their adaptation in different niches^3^.

Microbial social cheating can occur either at the genotypic or phenotypic level. The genotypic cheaters indicate the lost or mutation in specific gene(s) thus deficiency in the related biological function^14,27^; while the phenotypic cheaters are individuals with identical genetic background but silencing or down-regulation in public goods production (heterogeneous expression or division of labor)^25,28^. Despite the well-studied mechanisms of cheater control on the genotypic level^3^, those regarding to the phenotypic level remain largely unknown^3,25^; also unlike the definite significance of suppressing obligate genotypic cheaters^17^, the ecological roles of controlling phenotypic cheaters in mediating microbial population fitness have been rarely concerned^29^. Accordingly, lacking of the knowledge about phenotypic cheating limits our understanding of the cooperation behavior within microbial social communities.

Biofilms are extracellular matrix (ECM)-enclosed multicellular communities that sustain bacterial survival in diverse natural environments^30–32^, where the tightly associated cells are heterogeneously expressed with only a subpopulation of matrix producers^33–35^. As the ECM components (mainly include extracellular polysaccharides (EPS) and TasA fibers) are costly public goods shared by all cells within the biofilm, the nonproducing phenotypic cheaters can emerge, and thus disrupt the biofilm and community fitness^25,36^. Although a few studies have investigated the matrix production-cannibalism overlap and ECM privatization within biofilm individuals^25,29^, the molecular mechanism involved in punishment of nonproducing cheaters, as well as the ecological significance of the policing system in regulating population stability and fitness, remain unclear. *Bacillus velezensis* SQR9 (formerly *B*. *amyloliquefaciens* SQR9) is a well-studied beneficial rhizobacterium that form robust and highly structured biofilms on air-liquid interface and plant roots^37–40^. Production of toxic bacillunoic acids (BAs), encoded by a unique genomic island in strain SQR9, was proved to occur in subfraction of cells with the self-immunity ability induced by BAs during biofilm formation, where the nonproducing siblings will be lysed by BAs^41,42^. Based on the manifestation that the BA-mediated cannibalism enhanced biofilm formation of strain SQR9, we hypothesized the ECM and BAs synthesis can be co-regulated to restrain cheaters and sustain population stability. Using a combination of single-cell tracking technique, molecular approaches, and ecological evaluation, we demonstrated the ECM and BAs production are coordinated in the same subpopulation by the same regulator during biofilm formation, which enforces punishment of the nonproducing phenotypic cheaters to maintain community stabilization; also this genomic island-governed policing system is significant to promote community fitness in various conditions.

## Results

### Coordinated production of extracellular matrix (ECM) and autotoxin bacillunoic acids (BAs) punishes public goods-nonproducing cheaters in *B. velezensis* SQR9 community

To test the hypothesis that secretion of cannibal toxin eliminates the public goods-nonproducing cheaters in *B. velezensis* SQR9 community, we firstly tried to determine whether ECM (public goods) production and BAs (autotoxin) biosynthesis/BAs-induced self-immunity occur in the same subpopulation. We fused promoters for genes related to extracellular polysaccharides (EPS) and TasA fibers biosynthesis with *mCherry,* while the promoters for genes related to the autotoxin BAs biosynthesis and the self-immunity with *gfp,* obtained the *P_eps_-mCherry, P_tapA_-mCherry, P_bnaF_-gfp,* and *P_bnaAB_-gfp*, respectively. Their expression patterns were monitored using confocal laser scanning microscopy (CLSM) during the biofilm community formation. Photographs show that expression of the *P_eps_-mCherry, P_tapA_-mCherry, P_bnaF_-gfp*, and *P_bnaAB_-gfp* were all observed in a subpopulation cells of the whole community (Fig. 1), which suggests a differential expression pattern of each function among subpopulations during biofilm formation, where the ECM-nonproducers can be recognized as phenotypic cheaters^25^. Importantly, the overlay of the double fluorescent reporters indicates that ECM and BAs production is generally raised in the same subpopulation (Fig. 1; the yellow cells represent co-expression of *mCherry* and *gfp*); as expected, since the self-immunity gene *bnaAB* was reported to be specifically activated by endogenous BAs^42^, it was also preferentially expressed in the same subpopulation with ECM-producers (Fig. 1). These observations demonstrate a general coordination of ECM production and BAs synthesis/immunity in the same subpopulation of *B. velezensis* SQR9 biofilm community.

**Fig. 1.**
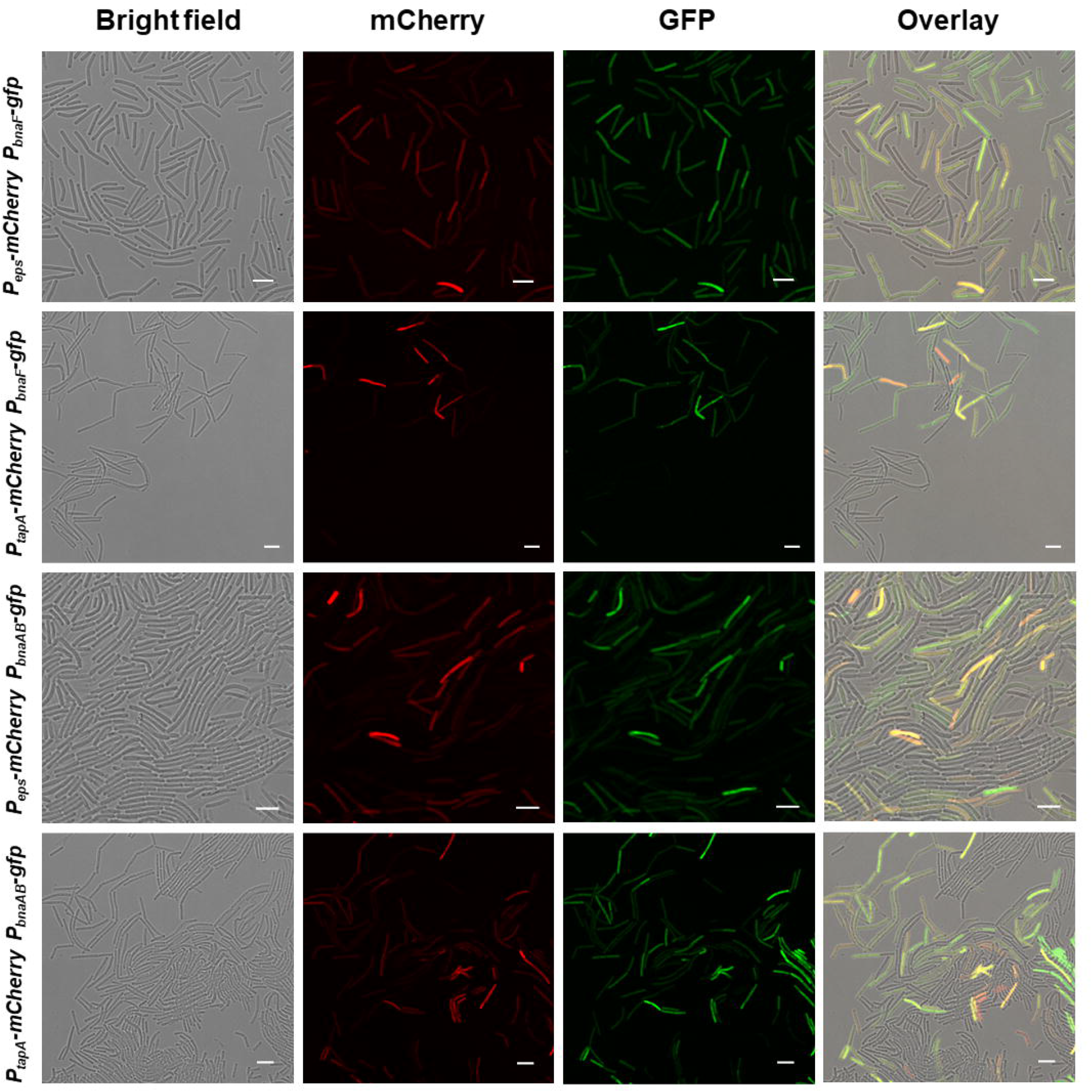
Expression of extracellular matrix (ECM) production and bacillunoic acids (BAs) biosynthesis/immunity were located in the same subpopulation. Colony cells of different double-labeled strains were visualized using a confocal laser scanning microscopy (CLSM) to monitor the distribution of fluorescence signal from different reporters. *P_eps_-mCherry* and *P_tapA_-mCherry* were used to indicate cells expressing extracellular polysaccharides (EPS) and TasA fibers production, respectively; *P_bnaF_-gfp* and *P_bnaAB_-gfp* were used to indicate cells expressing BAs synthesis and self-immunity, respectively. The bar represents 5 μm.

Based on the co-expression pattern, we postulated that the ECM-nonproducing cheaters, synchronously being sensitive to the BAs, will be killed by their siblings that produce both public goods ECM and the autotoxin BAs. Combining propidium iodide (a red-fluorescent dye for labeling dead cell) staining with reporter labelling, we monitored the cell death dynamics during the biofilm formation process in real time. It was observed that a portion of the cells that didn’t produce public ECM (Fig. 2A & 2B) or toxic BAs (Fig. 2C), or silenced in expression of the self-immunity gene *bnaAB* (Fig. 2D), were killed during the biofilm development process, while the corresponding producers remained alive throughout the incubation (red arrows indicate the dead cells in Fig. 2; Movies S1~S4). This lysis can be attributed to the BAs produced by the *gfp*-activated cells, as cannibalism of *B. velezensis* SQR9 was largely dependent on the production of this secondary metabolism^42^. Taken together, the double-labelling observation and cell death dynamics detection indicate that the subpopulation of ECM and BAs producers selectively punish the nonproducing siblings depend on a coordinately activated cell-differentiation pathway.

**Fig. 2.**
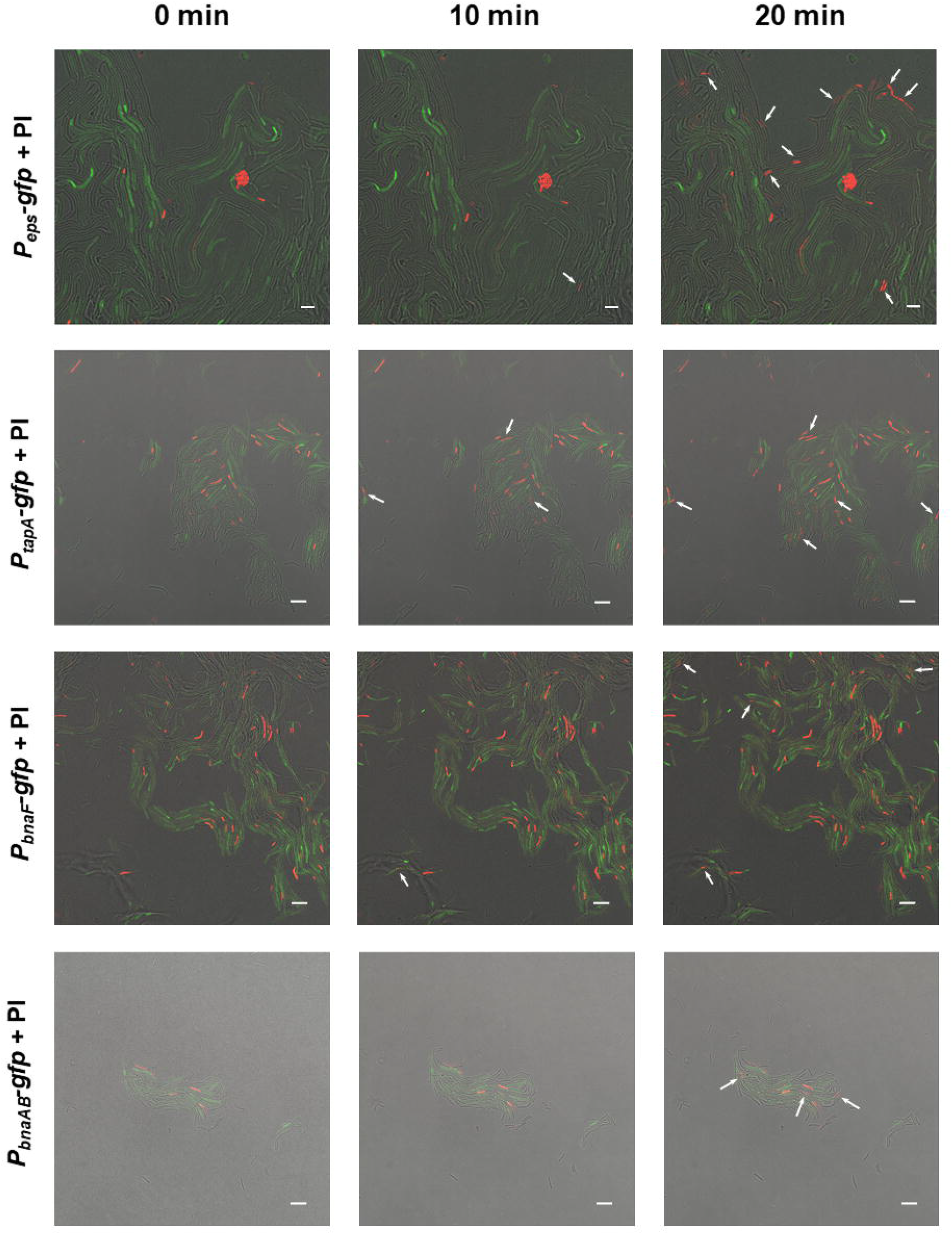
ECM and BAs producing subpopulation eliminated the nonproducing cheaters. Colony cells of different *gfp*-labeled strains were stained with propidium iodide (PI, a red-fluorescent dye for labeling dead cell) for 15 min, and then visualized by a CLSM to monitor the distribution of fluorescence signal from reporters and the PI dye, at 0, 10, and 20 min after treatment. *P_eps_-gfp* and *P_tapA_-gfp* were used to indicate cells expressing EPS and TasA fibers production, respectively; *P_bnaF_-gfp* and *P_bnaAB_-gfp* were used to indicate cells expressing BAs synthesis and self-immunity, respectively.

### Spo0A is the co-regulator for triggering ECM production and BAs synthesis/immunity

To identify the potential co-regulator(s) of ECM production and BAs synthesis/immunity in *B. velezensis* SQR9, we evaluated the BAs production in an array of mutants that known to be altered in ECM synthesis *(ΔdegU, ΔcomPA, ΔabrB, ΔsinI, ΔsinR,* and *Δspo0A),* by measuring their antagonism towards *B. velezensis* FZB42, a target strain specifically inhibited by BAs but no other antibiotics secreted by SQR9^41^. The crude extract of BAs of wild-type SQR9 showed remarkable antagonism to the of lawn of strain FZB42 (Fig. 3A & 3B); only *Δspo0A* but no other mutants (all with the equal cell density of the wild-type), revealed significantly reduced inhibition zone towards FZB42, and the complementary strain generally restored the antagonistic ability (Fig. 3A & 3B). Spo0A is a well-investigated master regulator that governs multiple physiological behaviors in *B. subtilis* and closely-related species^43,44^; as expected, the EPS production and biofilm formation was seriously impaired in *Δspo0A* (Fig. S1). Intriguingly, *Δspo0A* but neither its complementary strain nor the wild-type, can be substantially inhibited by the crude extracted BAs of strain SQR9, while *Δspo0A* was not inhibited by ΔGI3 that disabled in BAs production (Fig. 3C), suggesting Spo0A does participate in the immunity to BAs. In addition, we constructed *gfp* transcriptional fusions to the promoter of genes involved in ECM production (*eps* & *tapA)* and BAs biosynthesis/immunity *(bnaF/bnaAB),* and discovered that under both liquid culture (Fig. 3D) and plate colony conditions (Fig. S2), their expression level was significantly decreased in *Δspo0A* as compared with the wild-type, which was restored in the complementary strain *Δspo0A/spo0A.* These results suggest that the global regulator Spo0A is the co-regulator for controlling ECM production and BAs biosynthesis/immunity in *B. velezensis,* which is probably dependent on the transcriptional regulation of certain relevant genes.

**Fig. 3.**
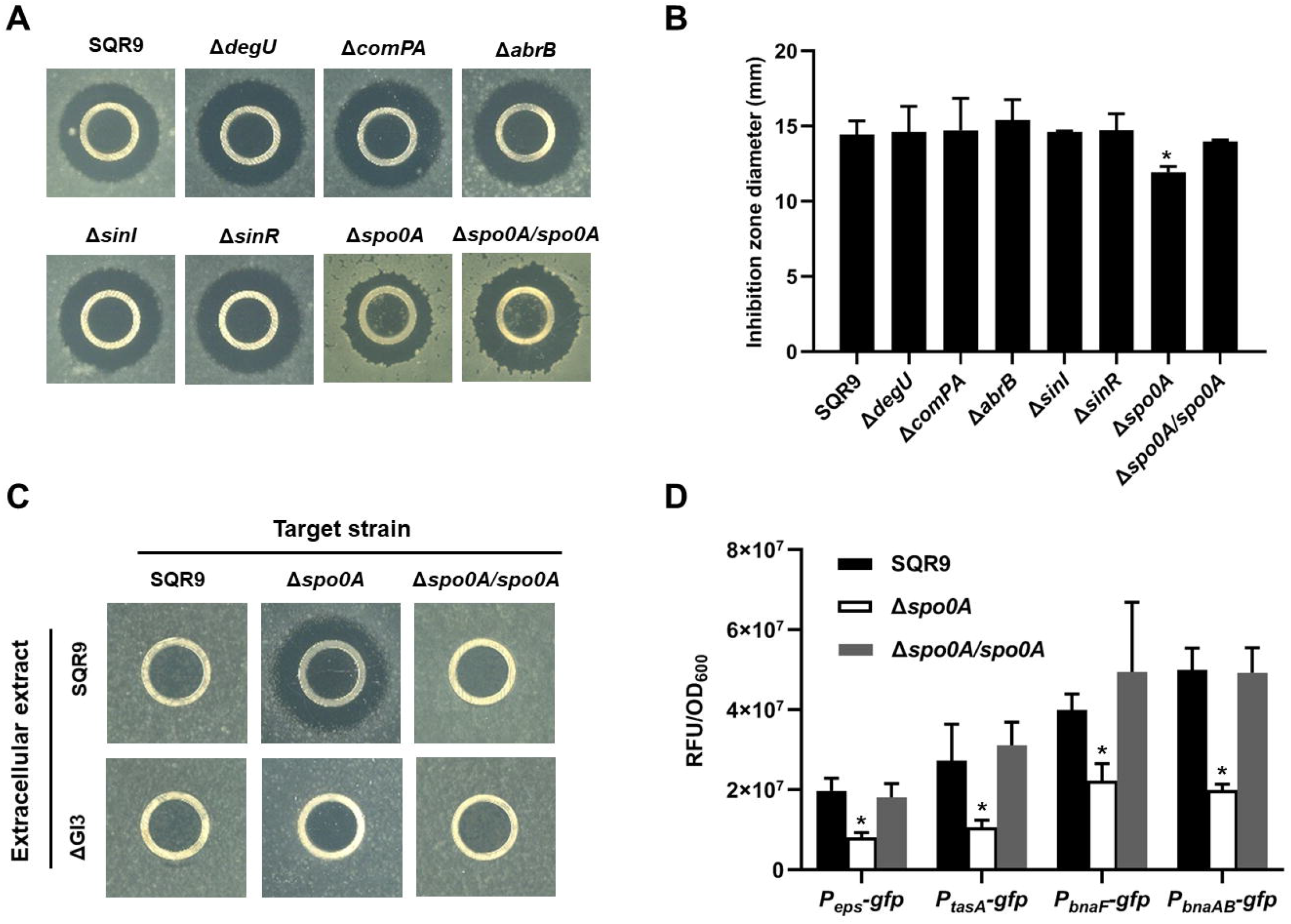
Spo0A is the co-regulator for triggering ECM production and BAs synthesis/immunity. (**A**) Inhibition of the lawn of *B. velezensis* FZB42 by the crude extracted BAs of wild-type SQR9, its different mutants altered in ECM production, and complementary strain *Δspo0A/spo0A.* (**B**) Diameter of the inhibition zones observed in (**A**). (**C**) Sensitivity of wild-type SQR9, *Δspo0A,* and *Δspo0A/spo0A* (as the lawn) to the extracellular extract of SQR9 and its mutant ΔGI3 that disable in BAs synthesis. (**D**) Expression level of *eps*, *tapA, bnaF,* and *bnaAB* in wild-type SQR9, *Δspo0A,* and *Δspo0A/spo0A,* as monitored by using *gfp* reporters fused to the corresponding promoters. Data are means and standard deviations from three biological replicates. * indicates significant difference with the Control (SQR9) column as analyzed by Duncan’s multiple range test (*P* < 0.05).

### Spo0A activates acetyl-CoA carboxylase (ACC) to support BAs synthesis and self-immunity

Despite the well-known Spo0A pathway in governing ECM production and biofilm formation in *Bacillus*^6^, how does Spo0A regulate BAs synthesis and self-immunity remains unknown. By using the biolayer interferometry analysis (BLI) for detecting molecular interaction, we revealed that the purified protein Spo0A cannot directly bind to the promoter of *bnaF,* suggesting it doesn’t induce BAs production through direct transcriptional activation (Fig. S3). Alternatively, Spo0A has been reported to stimulate the expression of *accDA* that encodes acetyl-CoA carboxylase^45,46^, which catalyzes acetyl-CoA to generate malonyl-CoA, an essential precursor for BAs biosynthesis (Fig. 4A)^41^; therefore we postulated *accDA* may be involved in the regulation of BAs production/immunity by Spo0A. We firstly verified the positive regulation of Spo0A on *accDA* expression in *B. velezensis* SQR9 by *gfp* fusion (Fig. 4B & Fig. S4). Since knockout of *accDA,* the essential gene for fatty acids biosynthesis, significantly impact bacterial growth, we alternatively constructed a strain in which the original promoter of *accDA* was replaced by a xylose-inducible promoter (*Pxyl*), and monitored its BAs synthesis/immunity under different xylose induction conditions. The SQR9-*Pxyl-accDA* lost the antagonism ability towards target strain FZB42 in the absence of xylose, while the inhibition was significantly enhanced with the induction of xylose in a dose-dependent manner (Fig. 4c & 4d). Since exogenous xylose didn’t influence the suppression of wild-type SQR9 on FZB42 (Fig. 4C & 4D), these results suggest that *accDA* expression positively contribute to BAs production. Importantly, the SQR9-*Pxyl-accDA* was proved to be sensitive to SQR9-produced BAs without xylose addition, and the immunity was gradually restored with xylose supplement (Fig. 4E). The xylose-induced transcription of *accDA,* also resulted in enhanced expression of genes involved in self-immunity *(bnaAB;* Fig. 4F & Fig. S5A) but not BAs synthesis *(bnaF;* Fig. 4F & Fig. S5B), as the AccDA-derived malonyl-CoA accumulation affects BAs production in a post-transcriptional manner. The CLSM photographs also reveal that the activation of *accDA (mCherry* fusion) and *bnaAB (gfp* fusion) was located in the same subpopulation cells (Fig. S6). Accordingly, these results indicate the positive regulation of Spo0A on BAs production/immunity in *B. velezensis* SQR9, is strongly dependent on *accDA* that encodes acetyl-CoA carboxylase.

**Fig. 4.**
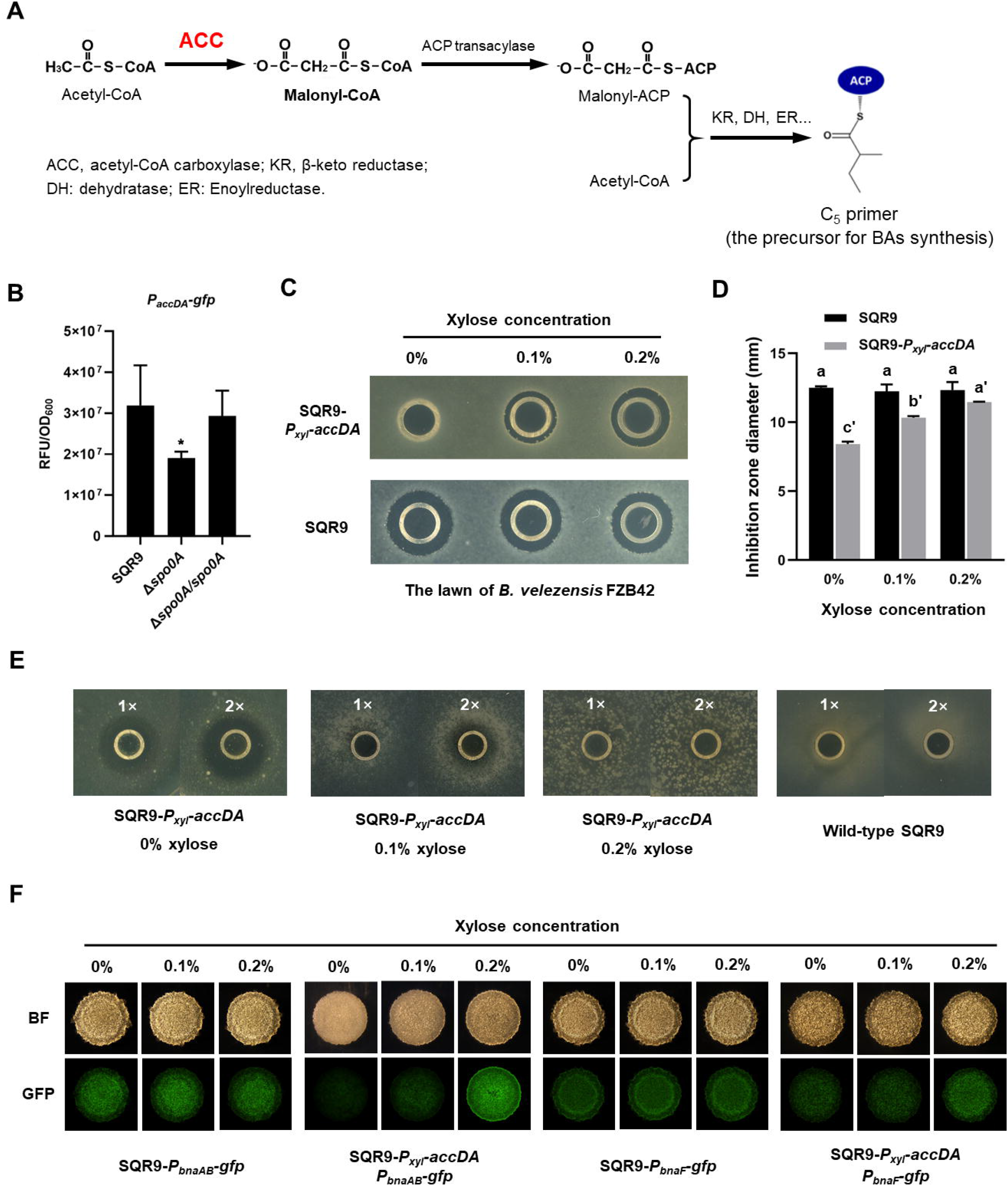
Spo0A activates acetyl-CoA carboxylase (ACC) for BAs synthesis and self-immunity. (**A**) Involvement of ACC in biosynthesis of BAs in *B. velezensis* SQR9. ACC catalyzes acetyl-CoA to generate malonyl-CoA, which is transformed to malonyl-ACP under the catalyzation of ACP transacylase; then malonyl-ACP and acetyl-CoA are aggregated into a C_5_ primer, the precursor for BAs synthesis. (**B**) Expression level of *accDA* in wild-type SQR9, *Δspo0A,* and *Δspo0A/spo0A,* as monitored by using the *P_accDA_-gfp* reporter. (**C**) Inhibition of the lawn of *B. velezensis* FZB42 by the crude extracted BAs of wild-type SQR9 and *SQR9-P_xyl_-accDA,* with addition of different concentrations of xylose (0%, 0.1% and 0.2%). (**D**) Diameter of the inhibition zones observed in (**C**). (**E**) Sensitivity of wild-type SQR9 and *SQR9-P_xyl_-accDA* (as the lawn) to the crude extracted BAs of SQR9 (100 μL (1×) or 200 μL (2×)), with addition of different concentrations of xylose (0%, 0.1%, and 0.2%). (**F**) Expression of *bnaF* and *bnaAB* in the colony cells of wild-type SQR9 and *SQR9-P_xyl_-accDA,* with addition of different concentrations of xylose (0%, 0.1% and 0.2%). Colonies were observed under both bright field (BF in the figure) and GFP channel, to monitor the florescence of *P_bnaF_-gfp* and *P_bnaAB_-gfp* reporters in different strains. Data are means and standard deviations from three biological replicates. * in (**B**) indicates significant difference (*P* <0.05) with the Control (SQR9) column as analyzed by Duncan’s multiple range tests; columns with different letters in (**D**) are statistically different according to the Duncan’s multiple range test (“a” for wild-type SQR9 under different concentrations of xylose and “a’“ for SQR9-*P_xyl_-accDA*; *P* <0.05).

### The co-regulation policing system enhances population stability and fitness

Having illustrated the molecular mechanism of the co-regulation pathway for punishing nonproducing cheaters in *B. velezensis* SQR9, we wondered the broad-spectrum ecological significance of this policing system for *B. velezensis* SQR9 in a community level. We constructed two mutants with disabled sanction mechanism, the *ΔbnaV* deficient in BAs synthesis (loss of the punishing weapon) and the SQR9*-P_43_-bnaAB* that continually expresses the self-immunity genes (cheaters cannot be punished by the weapon BAs), both mutants showed similar growth characteristics with the wild-type (Fig. S7). We firstly applied flow cytometry analysis to test whether lack of the policing system *(ΔbnaV* and *SQR9-P_43_-bnaAB)* impair the punishment of public goods nonproducing cheaters during biofilm formation. The proportion of matrix-producing cooperators (*eps* & *tapA* active cells) in the wild-type community, as well as the average expression level of corresponding genes, were significantly higher than that in the *ΔbnaV* or *SQR9-P_43_-bnaAB* community (Fig. 5A & 5B), suggesting the cheating individuals were not effectively controlled in the two mutants population. Consequently, the wild-type established a more vigorous biofilm as compared with the two mutants, as shown by the earlier initial progress, larger maximum biomass, and delayed dispersal process (prolonged stationary phase) (Fig. 5C & 5D). Additionally, the robust biofilm formed by the wild-type also endowed them stronger resistance against different stresses, including antibiotics, salinity, acid-base, and oxidation (Fig. 5D, Figs. S8 & S9).

**Fig. 5.**
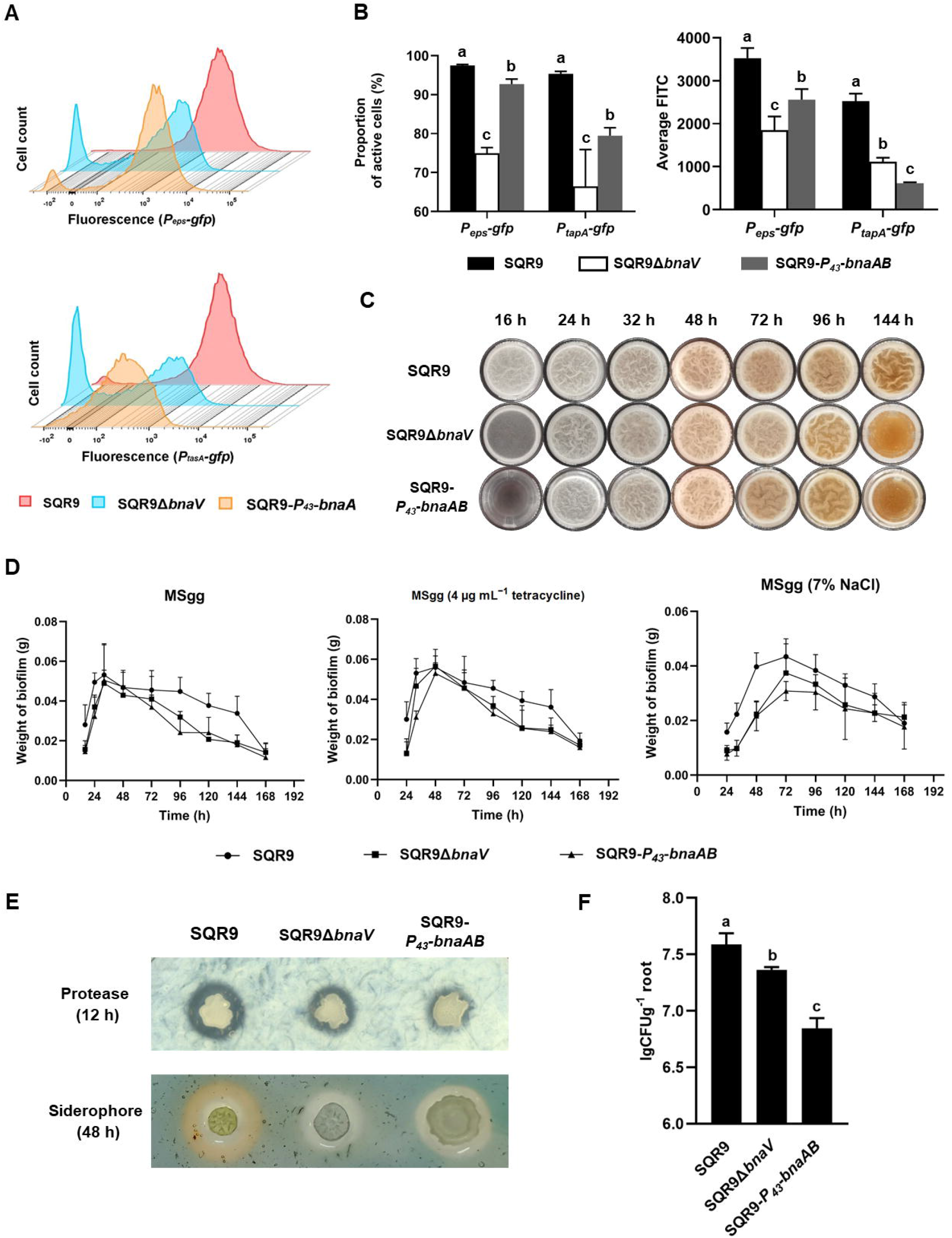
The co-regulation policing system eliminates cheaters and enhances population fitness. (**A**) Flow cytometry monitoring the expression of *P_eps_-gfp* and *P_tapA_-gfp* reporters in wild-type SQR9, *SQR9ΔbnaV* and *SQR9-P_43_-bnaAB.* (**B**) Proportion of the active cells (%) and average FITC in wild-type SQR9, SQR9Δ *bnaV* and SQR9 *-P_43_-bnaAB,* as monitored by *Peps-gfp* and *P_tapA_-gfp* reporters using flow cytometry. (**C**) Pellicle formation dynamics of wild-type SQR9, *SQR9ΔbnaV* and SQR9-*P_43_-bnaAB* in MSgg medium. (**D**) Pellicle weight dynamics of wild-type SQR9, *SQR9ΔbnaV*and *SQR9-P_43_-bnaAB* in MSgg medium under normal (corresponds to (**C**)) or stressed conditions (H_2_O_2_, tetacycline, or 7% NaCl). (**E**) Production of proteases and siderophore by wild-type SQR9, *SQR9ΔbnaV*and *SQR9-P_43_-bnaAB* colonies. (**F**) Comparison of root colonization of wild-type SQR9, *SQR9ΔbnaV* and *SQR9-P_43_-bnaAB.* Data are means and standard deviations from three biological replicates; columns with different letters are significantly different according to Duncan’s multiple range tests, *P* < 0.05.

Besides the well-known regulation on biofilm matrix production, Spo0A also controls the production of other public goods, such as proteases and siderophore^44,47^; it can be recognized as a critical switch that governs the cell transition from a free-living and fast-growing status (Spo0A-OFF), to a multicellular and cooperative style (Spo0A-ON)^34,48^. Intrinsically, the punishing targets of this policing system are supposed not limited to the matrix-nonproducing cheaters, but all of the Spo0A-OFF individuals (cells that don’t express the immune genes *bnaAB*). Therefore we determined the production of extracellular proteases and siderophore among the three strains, revealing that these public goods were also accumulated more in the wild-type than in these two mutants community (Fig. 5E & Fig. S10). Importantly, the wild-type SQR9 demonstrated a significantly stronger root colonization comparing with the two mutant strains losing the cheater punishing system (Fig. 5F). In summary, the Spo0A governed co-regulation punishment system effectively excludes the nonproducing cheaters of public goods in *B. velezensis* population, thereby improving the population stability and ecological fitness under different conditions.

## Discussion

Microbes have evolved diverse strategies for preventing cheaters in their communities. Despite the well-established mechanisms for controlling genotypic cheaters^3,16,17,49^, those regarding the phenotypic cheaters remain largely unknown^25,29^. Phenotypic cheaters are ordinarily derived from heterogeneous expression of different biological functions within a cell population^35^, this division of labor is postulated to afford bacterial community a better adaptation to unexpected environmental fluctuations^34^; however, when cells are supposed to be developed into a certain type in response to the surroundings, such as producing ECM to form surface-attached biofilm, or secreting extracellular enzymes to excavate resources, the individuals that don’t perform these assignments (but still share the community’s public goods) become actual cheaters and may disturb the community stability and fitness^15,29^. In the present study, we demonstrated that during biofilm formation, the beneficial rhizobacterium *B. velezensis* SQR9 engages a policing system that coordinately actives ECM production and autotoxin synthesis/immunity, to punish the phenotypic cheaters silencing in public goods secretion and reduce their proportion in the community (Fig. 6). This finding coincides with the coordinated cannibalism phenomenon reported in biofilm formation by *B. subtilis*^29^. Specifically, the toxic BAs for punishment is synthesized by a horizontal gene transfer (H GT)-acquired genomic island^41^, where its production is regulated by a precursor-dependent post-transcriptional manner (Fig. 4), and self-immunity is induced by the BAs through a two-component system^42^; importantly, this sanction mechanism not only facilitates ECM accumulation, but also contributes to enhanced production of other public goods including proteases and siderophore, thereby effectively improving the community fitness under different stressful conditions and in plant rhizosphere (Fig. 5). Actually, the coordination policing system eliminates phenotypic cheaters that stay in a fast-growing, motility phase (Spo0A~OFF state), to promote the population to a stationary, resource-mining phase (Spo0A-ON state) when environment required (Fig. 6). It should be noted that the phenotypic cheaters are not so obligate or detrimental, and this punishment is relatively temperate than those for genotypic cheaters^10,50^ as only a subpopulation of the cheaters were killed (Fig. 2); we think this scene is a balance between restraining the temporary cheaters and retaining the advantages of heterogeneous population^34,51^.

**Fig. 6.**
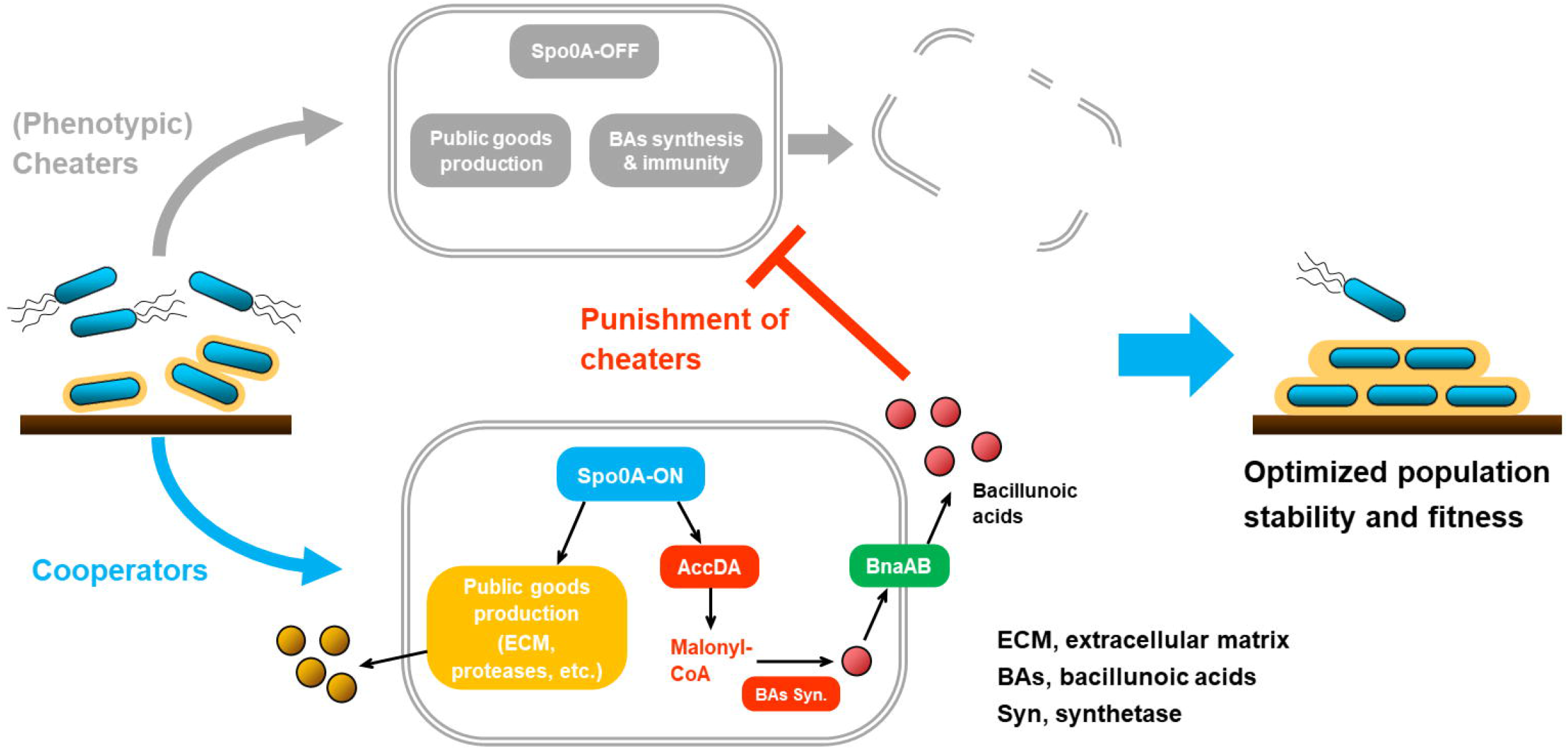
Working model and ecological significance of the co-regulation policing system in *B. velezensis.* In certain conditions (e.g., environmental or self-produced clues, surface attachments, etc.), *Bacillus* cells can differentiate into Spo0A-ON (~moderate phosphorylated) and Spo0A-OFF (unphosphorylated) subpopulation. The Spo0A-ON subpopulation are cooperators that produce public goods for the community, such as extracellular matrix (ECM) or proteases; simultaneously they express AccDA to produce malonyl-CoA as the precursor for bacillunoic acids (BAs) biosynthesis, and the endogenous autotoxin activates immunity-required transporter BnaAB to pump them out. Comparatively, the Spo0A-OFF subpopulation are phenotypic cheaters that silenced in public goods secretion, which are also disable in malonyl-CoA production and BAs biosynthesis/self-immunity. Consequently, the cooperators-produced BAs can effectively eliminate the cheating individuals, thereby enhancing the population stability and fitness.

The diversified cheater-controlling mechanisms used by microorganisms, reveal different applicability features and can occur in various types of microbial cooperation^17^. For instance, kin discrimination is effective for controlling non-kin cheaters with different genetic backgrounds^52,53^, but appears incapable of preventing spontaneous genotypic cheaters in the same population, quite apart from the phenotypic cheaters^3^; facultative cooperation enables microbial population to optimize the occasion for producing public goods, which is an economic-style strategy for minimizing resource exploitation by cheaters while is unlikely to suppress them directly^20,54,55^; partial privatization and spatial structuring can immediately restrain the cheaters by physical separation^25,26^. Comparatively, the targeted benefit (private benefit) and punishment mechanisms afford cooperators direct fitness advantage over cheaters^10,22^, especially the latter precisely antagonizes the cheating individuals to eliminate them from the community^4,10^. The punishment strategy is usually elaborately regulated by QS or QS-like system for coupling the public goods production and autotoxins synthesis/immunity^56^, therefore it is both complicated for cheaters to overcome and costly for cooperators to implement^16^. Here we prove that the policing system in *B. velezensis* SQR9 contributes to optimized cell differentiation and population fitness, suggesting its ecological benefits does overcome the costs for expressing antibiotic production and immunity (Fig. 5). Alternatively, this sanction system can work in concert with privatization strategy to collectively prevent cheater invasion during biofilm formation^25^.

Interestingly, the secondary metabolites applied by *B. velezensis* SQR9 to punish the cheaters, are governed by a unique genomic island acquired through HGT^41^. Since *accDA* that is important for both BAs biosynthesis and the corresponding self-immunity, and *eps* and *tapA* operon required for ECM production, are all activated by Spo0A with moderate phosphorylation level (Fig. 4)^6,42,45^, these genes constitute an ingenious co-regulatory network to appoint the cooperators to be BAs producers and defenders, while the cheaters to be sensitive individuals that can be eliminated (Figs. 1 & 2). It was known that clusters carrying antibiotic biosynthesis and resistance genes (ARGs) are usually transformed among microbes through HGT in natural environment^57,58^, but since these elements also brought certain costs such as DNA replication and metabolic burden, they must produce considerable benefits to be reserved in the new host. Here the SQR9-acquired GI3 not only act as a weapon for antagonizing closely related competitors^41^, but also establishes a policing system for punishing cheaters within the internal community. We consider this dual function of the antibacterial fatty acids could explain why this large cluster was integrated in the genome of *B. velezensis* SQR9, and this case can provide inspirations for discovering novel molecular regulatory mechanisms and understanding microbial evolution events.

In conclusion, the present study highlights the beneficial rhizobacterium *B. velezensis* SQR9 engages a policing system that coordinately actives ECM production and autotoxin synthesis/immunity, to eliminate the phenotypic cheaters silencing in public goods secretion thereby enhancing the community fitness. This study provides insights of the molecular mechanism involved in controlling phenotypic cheaters, as well as the ecological roles of the policing system, which deepens our understanding of maintenance and evolution of microbial cooperation.

## Materials and Methods

### Bacterial strains and growth conditions

The strains and plasmids used in this study are listed in Table S1. *Bacillus velezensis* SQR9 (formerly *B. amyloliquefaciens* SQR9, China General Microbiology Culture Collection Center (CGMCC) accession no. 5808) was used throughout this study. *B. velezensis* FZB42 (Bacillus Genetic Stock Center (BGSC) accession no. 10A6) was used to test the bacillunoic acids (BAs) production by wild-type SQR9 and its mutants. *Escherichia coli* TOP 10 (Invitrogen, Shanghai, China) was used as the host for all plasmids. *E. coli* BL21 (DE3) (Invitrogen, Shanghai, China) was used as the host for recombinant protein expression. All strains were routinely grown at 37°C in low-salt Luria-Bertani (LLB) medium (10 g L^-1^ peptone, 5 g L^-1^ yeast extract, 3 g L^-1^ NaCl). For biofilm formation, *B. velezensis* SQR9 and its mutants were cultivated in MSgg medium (5 mM potassium phosphate, 100 mM morpholine propanesulfonic acid, 2 mM MgCl2, 700 μM CaCl2, 50 μM MnCl_2_, 50 μM FeCl_3_, 1 μM ZnCl_2_, 2 mM thiamine, 0.5% glycerol, 0.5% glutamate, 50 μg of tryptophan per milliliter, 50 μg of phenylalanine per milliliter, and 50 μg of threonineper milliliter) at 37°C^59^. To collect the fermentation supernatant for antagonism assessment, *B. velezensis* SQR9 and its mutants were cultured in Landy medium^60^ containing 20 g L^-1^ glucose and 1 g L^-1^ yeast extract. When necessary, antibiotics were added to the medium at the following final concentrations: zeocin, 20 μg mL^-1^; spectinomycin, 100 μg mL^-1^; kanamycin, 30 μg mL^-1^; ampicillin, 100 μg mL^-1^; chloramphenicol, 5 μg mL^-1^ for *B. velezensis* strains and 12.5 μg mL^-1^ for *E. coli* strains; erythromycin, 1 μg mL^-1^ for *B. velezensis* strains and 200 μg mL^-1^ for *E. coli* strains. The medium was solidified with 2% agar.

### Reporter construction

For single-labelled strain, the promoter region of the testing gene and *gfp* fragment were fused through overlap PCR, and this transcriptional fusion was cloned into vector pNW33n using primers listed in Table S2. For double-labelled strains, one promoter region was fused with *gfp* fragment and the other promoter region was fused with *mCherry* fragment. The two fusions were then fused in opposite transcription directions and cloned into vector pNW33n using primers listed in Table S2. All constructions were transferred into competent cells of *B. velezensis* SQR9 and mutants when required.

### Promoter replacement

Strain SQR9-*P_xyl_-accDA* was constructed by replacing the original promoter of *accDA (P_accDA_*) by a xylose-inducible promoter *P_xyl_.* The approximately 800 bp fragments of upstream and downstream of the *P_accDA_* region were amplified from the genomic DNA of strain SQR9; the Spc^r^ fragment was amplified from plasmid P7S6^61^, and the *Pxyl* promoter was amplified from the plasmid PWH1510^62^. The four fragments were fused using overlap PCR in the order of the upstream fragment, Spc^r^, *P_xyl_*, and the downstream fragment. The fusion was transferred into competent cells of *B. velezensis* SQR9 for generating transformants. Strain SQR9-*P_43_-bnaAB* was obtained by replacing the original promoter *(P_bnaAB_*) by a constitutive promoter *P_43_*. The primers used for constructing the four-fragment fusion are listed in Table S2.

### Fluorescence microscopy

Cells were inoculated from a fresh pre-culture and grown to mid-exponential growth at 37°C in LLB medium. Bacterial cultures were centrifuged at 4000 × g for 5 min, the pellets were washed and suspended in liquid MSgg to reach an OD_600_ of 1.0. One μL suspension was placed on solid MSgg medium and were cultured at 37°C for 12 h. Agarose MSgg pads were then inverted on a glass bottom dish (Nest). Cells were imaged using the Leica TCS SP8 microscope with the 63 × oil-immersion objective lens. For GFP observation, the excitation wavelength was 488 nm and the emission wavelength was 500~560 nm; for mCherry observation, the excitation wavelength was 587 nm and the emission wavelength was 590~630 nm. Wild-type biofilms containing no fluorescent fusions were analyzed to determine the background fluorescence.

For time-lapse experiment, after staining with propidium iodide (PI) for 15 min, images of colonies on the agarose pad were recorded for 20 min, with interval of 5 min. Image acquisitions were also performed with the Leica TCS SP8 microscope with the 63 × oil-immersion objective lens. Detectors and filter set for monitoring of GFP and PI (excitation wavelength of 536 nm and emission wavelength of 608~652 nm) were used.

### Preparation of the crude extract of BAs

The crude extract of BAs was prepared by thin layer chromatography (TLC). According to a previous study^41^, the fermentation supernatant of strain SQR9 were separated on a TLC plate, and the inhibition zone on the lawn of strain FZB42 indicated the position of BAs. Then, silica gel powder with BAs was scraped and extracted by MeOH, which was used as the crude extract of BAs.

### Oxford cup assay

Inhibition of different SQR9-derived mutants on *B. velezensis* FZB42 was evaluated by Oxford cup method. The suspension of strain FZB42 (~10^6^ CFU mL^-1^) was spread onto LLB plates (10 × 10 cm) to grow as a bacterial lawn. A volume of 100 μL crude extract of BAs produced by different mutants was injected into an Oxford cup on the lawn of strain FZB42. The plates were placed at 22°C until a clear zone formed around the cup, and the inhibition diameter was scored. Each treatment includes three biological replicates.

### BAs-sensitivity assessment

Cells were inoculated from a fresh pre-culture and grown to mid-exponential growth at 37°C in LLB medium. Afterwards, diluted cell suspension (~10^6^ CFU mL^-1^) was spread onto LLB plates to grow as a bacterial lawn. A volume of 100 μL crude extract of BAs from the wild-type SQR9 was injected into an Oxford cup on the lawn. The plates were placed at 22°C for observation and determination of the inhibition zone. Each treatment includes three biological replicates.

### Biolayer interferometry (BLI) measurements

To confirm whether Spo0A can bind *P_bnaF_* directly, determination of binding kinetics was performed on an Octet® RED96 device (ForteBio, Inc., Menlo Park, US) at 25°C with orbital sensor agitation at 1000 rpm. Streptavidin (SA) sensor tips (ForteBio) were used to immobilize 100 nM biotin-labeled *P_bnaF_*. Then, a baseline measurement was performed in the buffer PBST (PBS, 0.1% BSA, 0.02% Tween-20) for 300 s. The binding of Spo0A at different concentrations (100 nM, 250 nM, 500 nM, and 1000 nM) to *P_bnaF_* was recorded for 600 s followed by monitoring protein dissociation using PBST for another 600 s. The BLI data for each binding event were summarized as a “nm shift” (the wavelength/spectral shift in nanometers) and KD values determined by fitting to a 1:1 binding model.

### Promoter activity testing via fluorescence intensity

For colony fluorescence, cells were inoculated from a pre-culture into fresh LLB medium and grown at 37°C with 170 rpm shaking until OD_600_ reached 0.5. One μL of the suspension were inoculated on solid LLB medium and were cultured at 37°C. Colony morphology and fluorescence were recorded by the stereoscope. ImageJ software was used to measure GFP intensity. For liquid culture fluorescence, overnight cultures were transferred to fresh LLB medium. Fluorescence intensity was determined by a microtiter plate reader. Each treatment includes three biological replicates.

### Xylose induction assay

For the xylose-induced BAs production assay, 30 μL overnight culture of SQR9-*P_xyl_-accDA* or wild-type SQR9 was transferred respectively into 3 mL fresh LLB liquid with different concentrations of xylose (0%, 0.1%, 0.2%) and incubated at 37°C, 170 rpm for 24 h. Cell suspensions were adjusted to the same OD_600_ and were centrifuged at 12000 × g for 1 min. The cell-free supernatant was mixed with MeOH (volume ratio 2:1) to extract BAs. A volume of 100 μL extract was injected into an Oxford cup on the lawn of strain FZB42 (as described above). The plates were placed at 22°C.

For the xylose-induced self-immunity assay, strain SQR9-*P_xyl_-accDA* was grown in LLB without xylose for 24 h. Cell suspension was spread onto LLB plates containing different concentrations of xylose (0%, 0.1% and 0.2%) to grow as the lawn. A volume of 100 μL (1×) or 200 μL (2×) crude extract of BAs from the wild-type SQR9 was injected into an Oxford cup on the lawn. The plates were placed at 22°C.

For xylose-induced gene expression assay, cells were inoculated from a pre-culture into fresh LLB medium with different concentrations of xylose (0%, 0.1%, 0.2%), and were grown at 37°C with 170 rpm shaking until OD_600_ reached 0.5. One μL of suspension was inoculated on solid LLB medium and was cultured at 37°C, colony morphology and fluorescence were recorded by the stereoscope.

Each treatment in these assays includes three biological replicates.

### Biofilm formation

Cells were inoculated from a fresh pre-culture and grown to mid-exponential growth at 37°C in LLB medium. Bacterial cultures were centrifuged at 4000 × g for 5 min, the pellets were washed and suspended in MSgg medium to an OD_600_ of 1.0. For colony observation, 1 μL of suspension were inoculated on solid MSgg medium and were cultured at 37°C, then the colony morphology was recorded by the stereoscope. For pellicle observation, suspension was inoculated into MSgg medium with a final concentration of 1% in a microtiter plate well, and the cultures were incubated at 37°C without shaking.

Besides, the ability of strain to form biofilm under stress was measured in the 48-well microtiter plate according to the method described above. When required, reagents that simulate stress were supplemented in the MSgg medium before inoculating, including oxidative stress (0.0025% H_2_O_2_), salt stress (7% NaCl), acid stress (pH 5), alkaline stress (pH 8), and antibiotic stress (4 μg mL^-1^ tetracycline or 20 μg mL^-1^ streptomycin). The amount of reagent added was determined according to a concentration gradient in pre-experiment, and a concentration was chosen to inhibit wild-type growth without killing it. At different stages of biofilm development (initiation, progress, maturity, and dispersal), the MSgg medium underneath the biofilm was carefully removed by pipetting and then the biofilm was taken and weighed.

Each treatment includes three biological replicates.

### Flow cytometry

Biofilms were collected and re-suspended in 1 mL PBS buffer, and single cells were obtained after mild sonication. Cells were centrifuged at 4000 × g for 5 min and washed briefly with PBS. For flow cytometry, cells were diluted to 1:100 in PBS and measured on BD FACSCanto II. For GFP fluorescence, the laser excitation was 488 nm and coupled with 500-560 nm. Every sample was analyzed for 20000 events. FlowJo V10 software was used for data analysis and graphs creating. Three replicates for each treatment were analyzed.

### Root colonization assay in hydroponic culture

Bacterial suspension was inoculated into 1/4 Murashige-Skoog medium to make the final OD_600_ value to be 0.1, into which sterile cucumber seedlings with three true leaves were immersed. After cultured with slowly shaking for two days, cells colonized on cucumber roots were determined by plate colony counting. In detail, roots were washed eight times in PBS to remove free and weakly attached bacterial cells. After vortexing for 5 min until colonized bacteria were detached from roots, 100 μL of the bacterial suspension was plated onto LLB agar plates for quantification. Each treatment includes three biological replicates.

### Measurement of public goods production

Qualitative measurement of proteases production was done by inoculating 1 μL of bacterial suspension on solid 2% skim milk medium and cultured at 30°C until transparent zone formed around colonies; quantitative measurements of alkaline protease and neutral protease activity were conducted according to a previous study^63^. Qualitative and quantitative measurement of siderophore production were based on the universal chemical assay described by Schwyn and Neilands^64^. Each treatment includes three biological replicates.

## Acknowledgements

This work was financially supported by the National Natural Science Foundation of China (31870096, 42090064, 31972512, 32072665, and 32072675), the Fundamental Research Funds for the Central Universities (KYZZ2022003), and the National Key Research and Development Program (2021YFD1900300).

