## Supplementary Materials for "A toxin-mediated policing system in *Bacillus* improves population fitness via penalizing non-cooperating phenotypic cheaters"

**This file includes:**

Figs. S1 to S10

Tables S1 to S2

Legends for movies S1 to S4

**Other Supplementary Materials for this manuscript includes the following:**

Movies S1 to S4

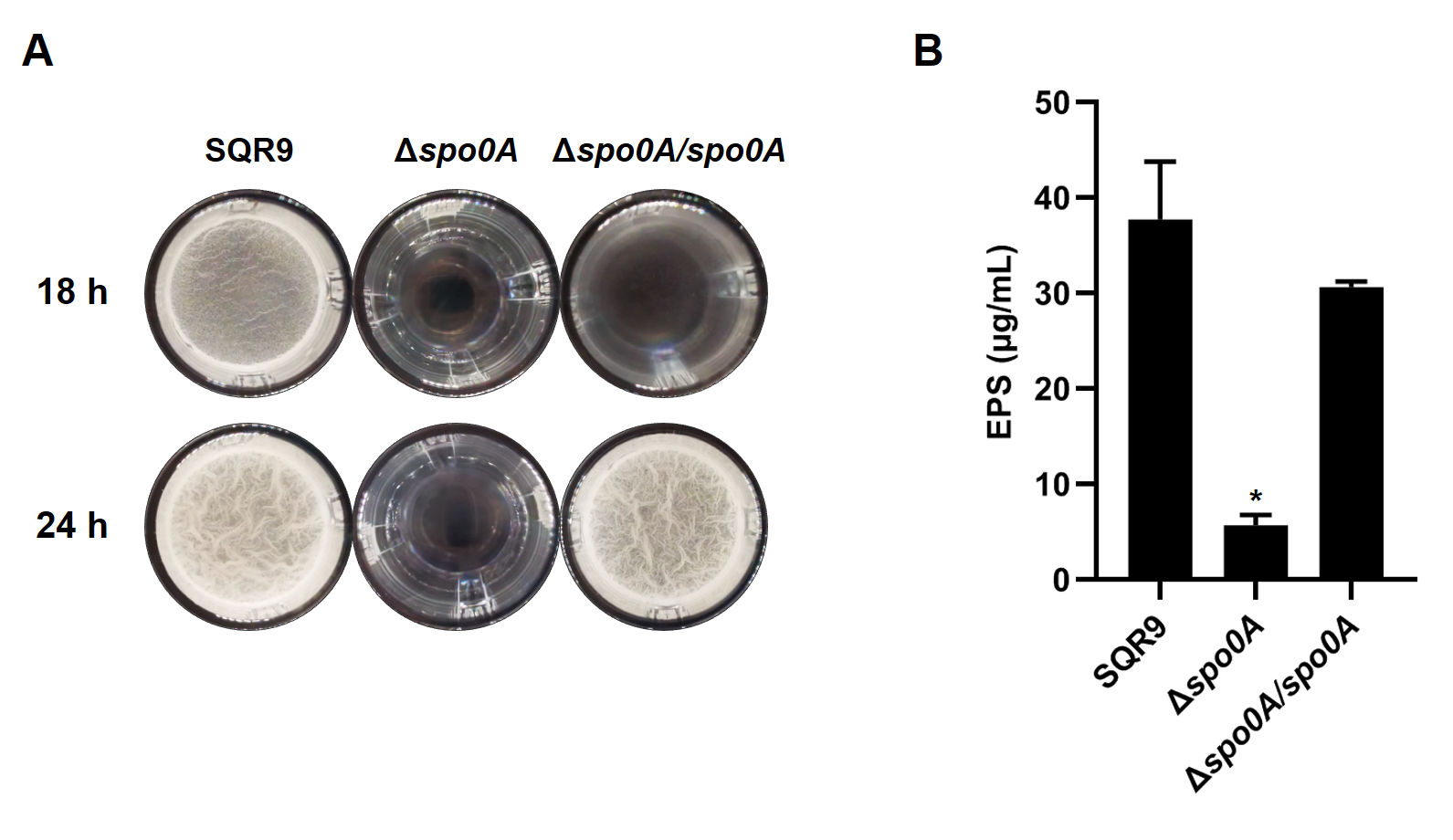

**Fig. S1. Pellicle formation (A) and extracellular polysaccharides (EPS) production (B) by wild-type SQR9, Δ*spo0A*, and Δ*spo0A/spo0A*.** ^*^ indicates significant difference (*P* < 0.05) with the Control (SQR9) column as analyzed by Duncan’s multiple range test (**B**).

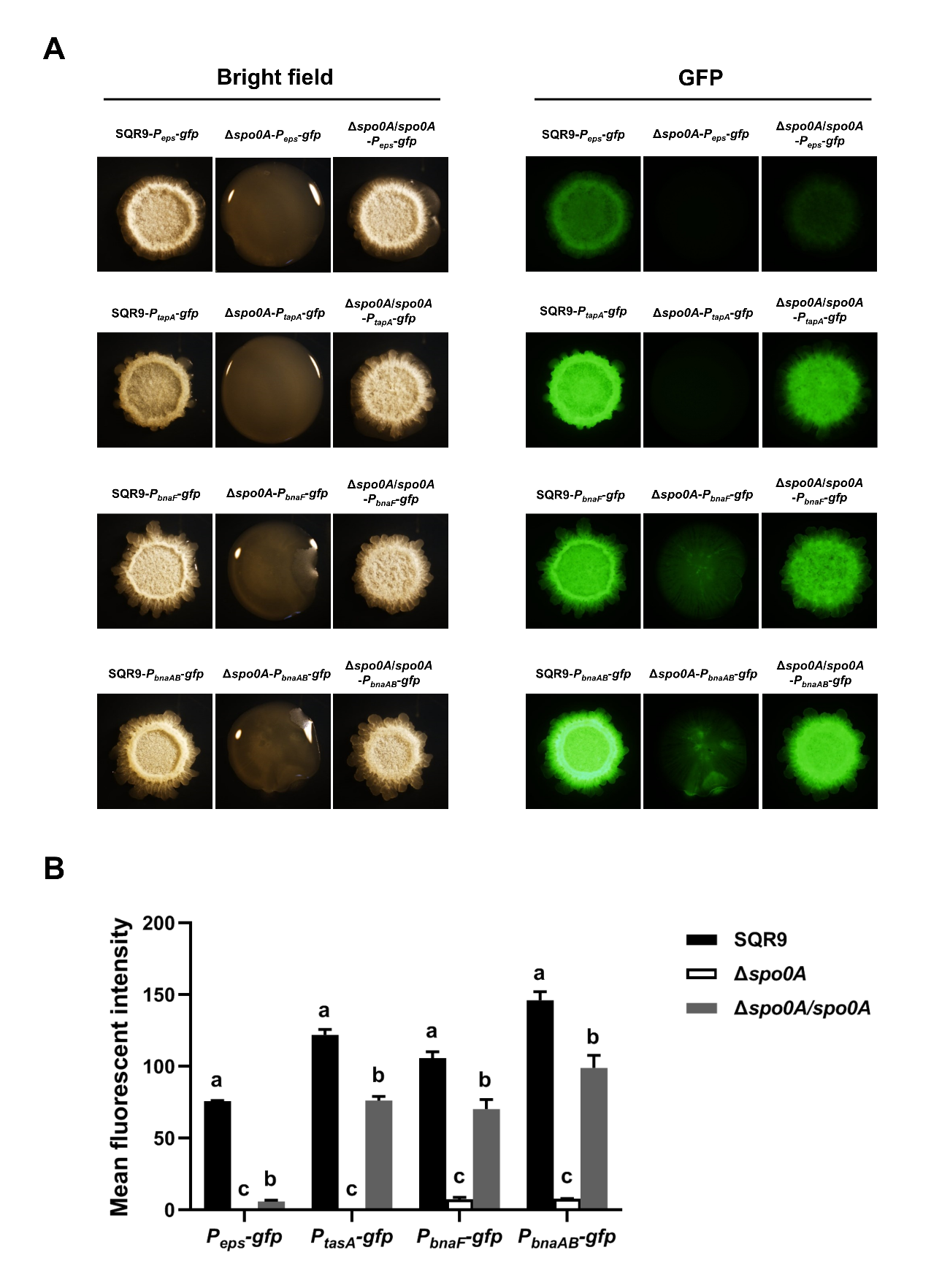

**Fig. S2. Expression level of *eps*, *tapA*, *bnaF*, and *bnaAB* in the colony cells of wild-type SQR9, Δ*spo0A*, and Δ*spo0A/spo0A*, as monitored by using *gfp* reporters fused to corresponding promoters.** (**A**) Colonies were observed under both bright field and GFP channel, to monitor the florescence of *P_eps_-gfp*, *P_tapA_-gfp*, *P_bnaF_-gfp*, and *P_bnaAB_-gfp* reporters in different strains. **(B**) The mean florescent intensity of different *gfp* reporters as observed in (**A**). Data are means and standard deviations from three biological replicates. Columns with different letters in a same group are statistically different according to the Duncan’s multiple range test (*P* < 0.05).

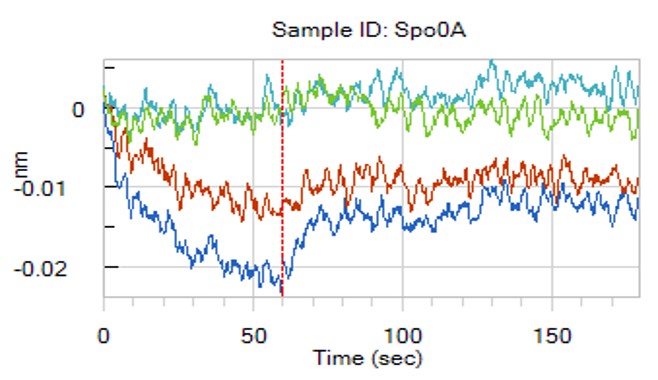

**Fig. S3. Interaction between the purified protein Spo0A and the promoter of *bnaF* (*P_bnaF_*) as determined by Biolayer interferometry data (BLI).**

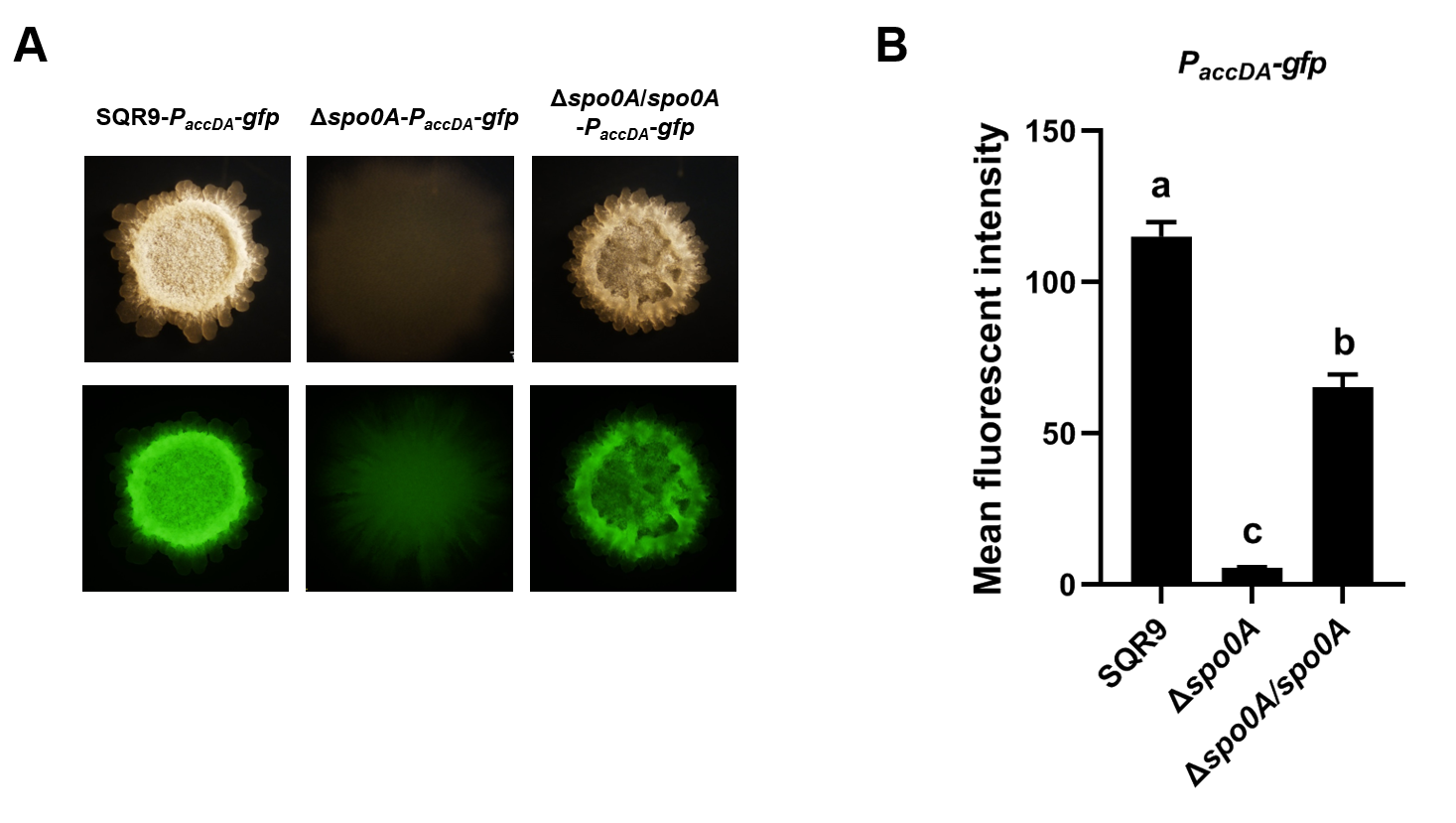
 **Fig. S4. Expression of *accDA* in the colony cells of wild-type SQR9, Δ*spo0A*, and Δ*spo0A/spo0A*, as monitored by using *gfp* reporters fused to promoter of *accDA*.** (**A**) Colonies were observed under both bright field and GFP channel, to monitor the florescence of *P_accDA_-gfp* in different strains. (**B**) The mean florescent intensity of different *gfp* reporters as observed in (**A**). Data are means and standard deviations from three biological replicates. Columns with different letters are statistically different according to the Duncan’s multiple range test (*P* < 0.05).

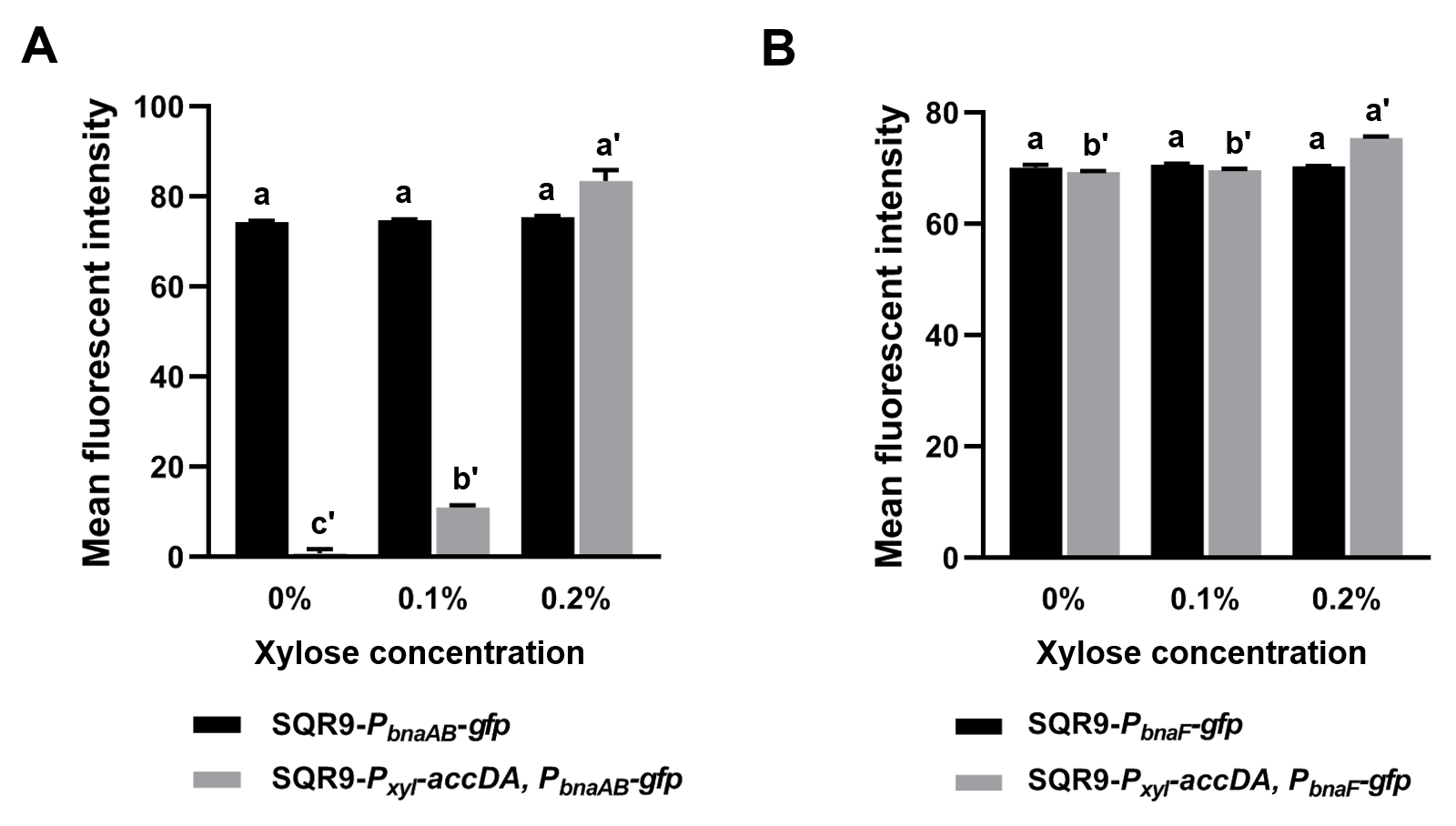

**Fig. S5. Expression level of *bnaAB* (A) and *bnaF* (B) in the colony cells of wild-type SQR9 and SQR9-*P_xyl_*-*accDA*, with addition of different concentrations of xylose (0%, 0.1%, and 0.2%).** The mean florescent intensity of different *gfp* reporters corresponds to the colonies shown in **Fig. 4F**. Data are means and standard deviations from three biological replicates. Columns with different letters are statistically different according to the Duncan’s multiple range test (“a” for SQR9-*P_bnaAB_*-*gfp* or SQR9-*P_bnaF_*-*gfp* under different concentrations of xylose, and “a'” for SQR9-*P_xyl_*-*accDA*, *P_bnaAB_*-*gfp* or SQR9-*P_xyl_*-*accDA*, *P_bnaF_*-*gfp*; *P* < 0.05).

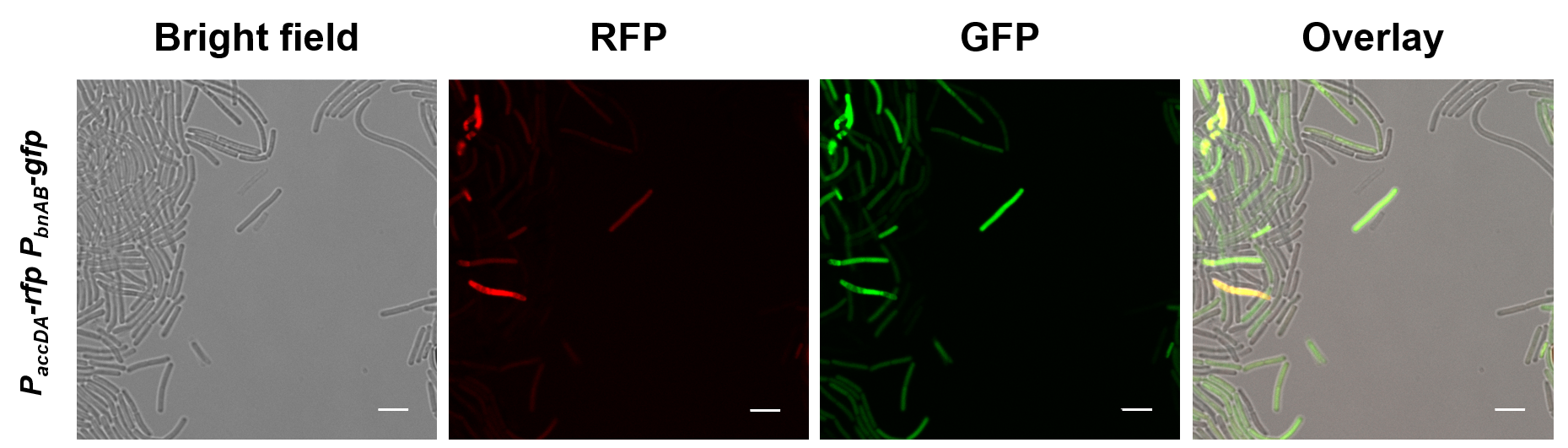
 **Fig. S6. Production of acetyl-CoA carboxylase and bacillunoic acids (BAs) immunity were located in the same subpopulation.** Colony cells of different double-labeled strains were visualized using a confocal laser scanning microscopy (CLSM) to monitor the distribution of fluorescence signal from different reporters. *P_accDA_-mCherry* and *P_bnaAB_-gfp* were used to indicate cells expressing acetyl-CoA carboxylase and BAs self-immunity, respectively. The bar represents 5 μm.

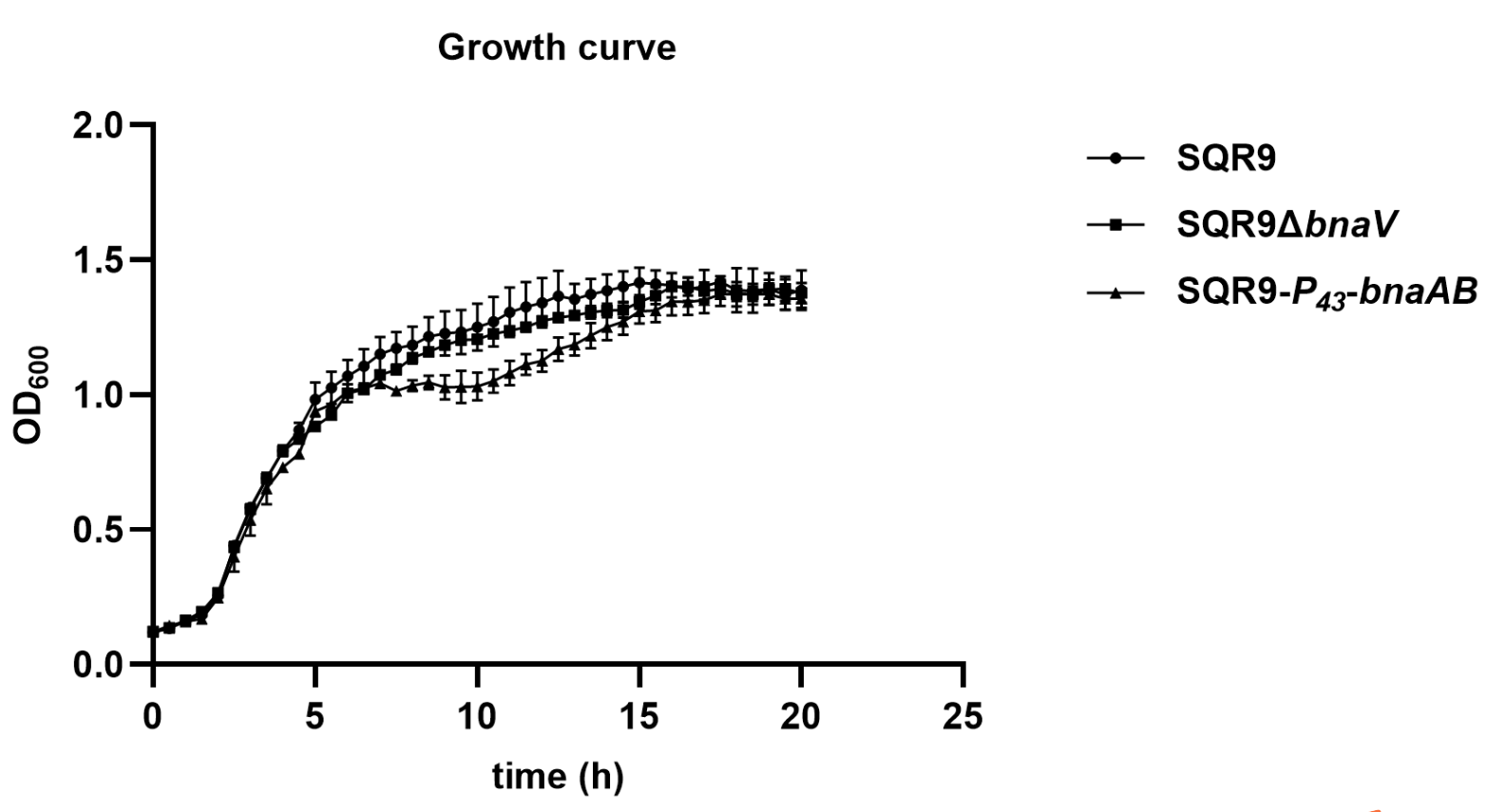

**Fig. S7. Growth curves of wild-type SQR9, SQR9Δ*bnaV*, and** **SQR9-*P_43_-bnaAB*.**

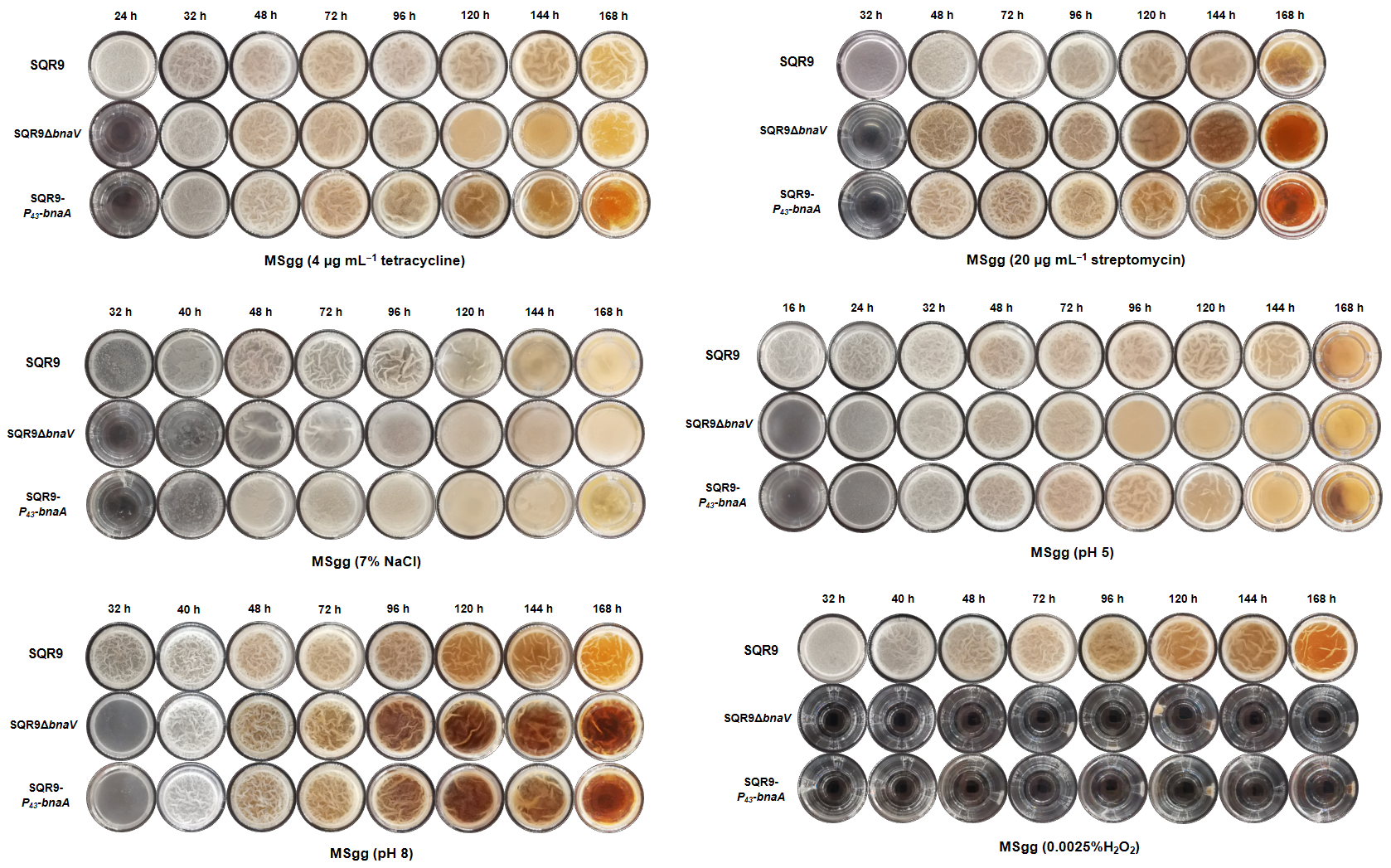

**Fig. S8. Dynamic pellicle formation of wild-type SQR9, SQR9Δ*bnaV*, and SQR9-*P_43_-bnaAB* in MSgg medium under different stressed conditions.**

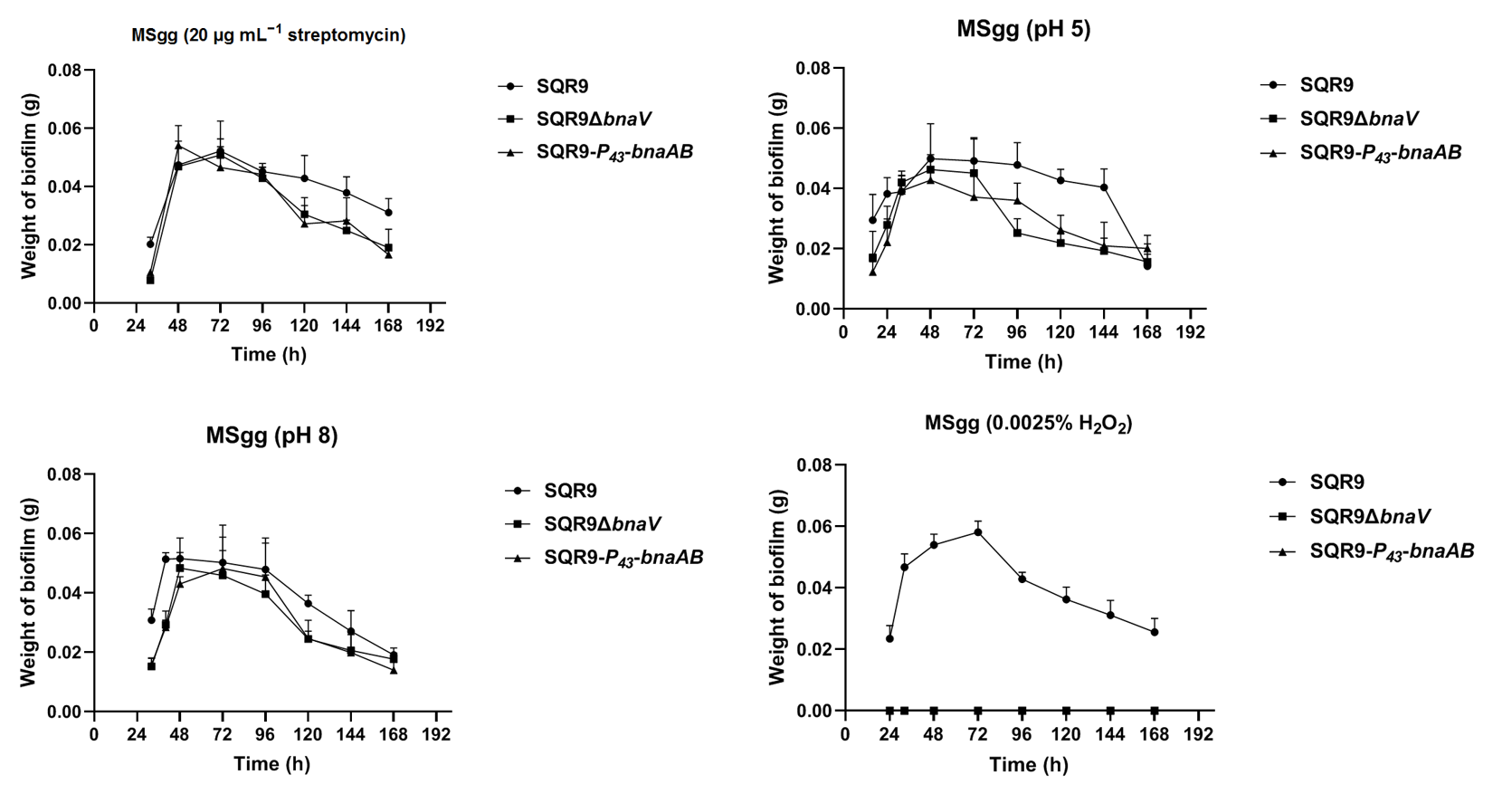

**Fig. S9.** **Dynamic pellicle weight of wild-type SQR9, SQR9Δ*bnaV*, and SQR9-*P_43_-bnaAB* in MSgg medium under different stressed conditions.**

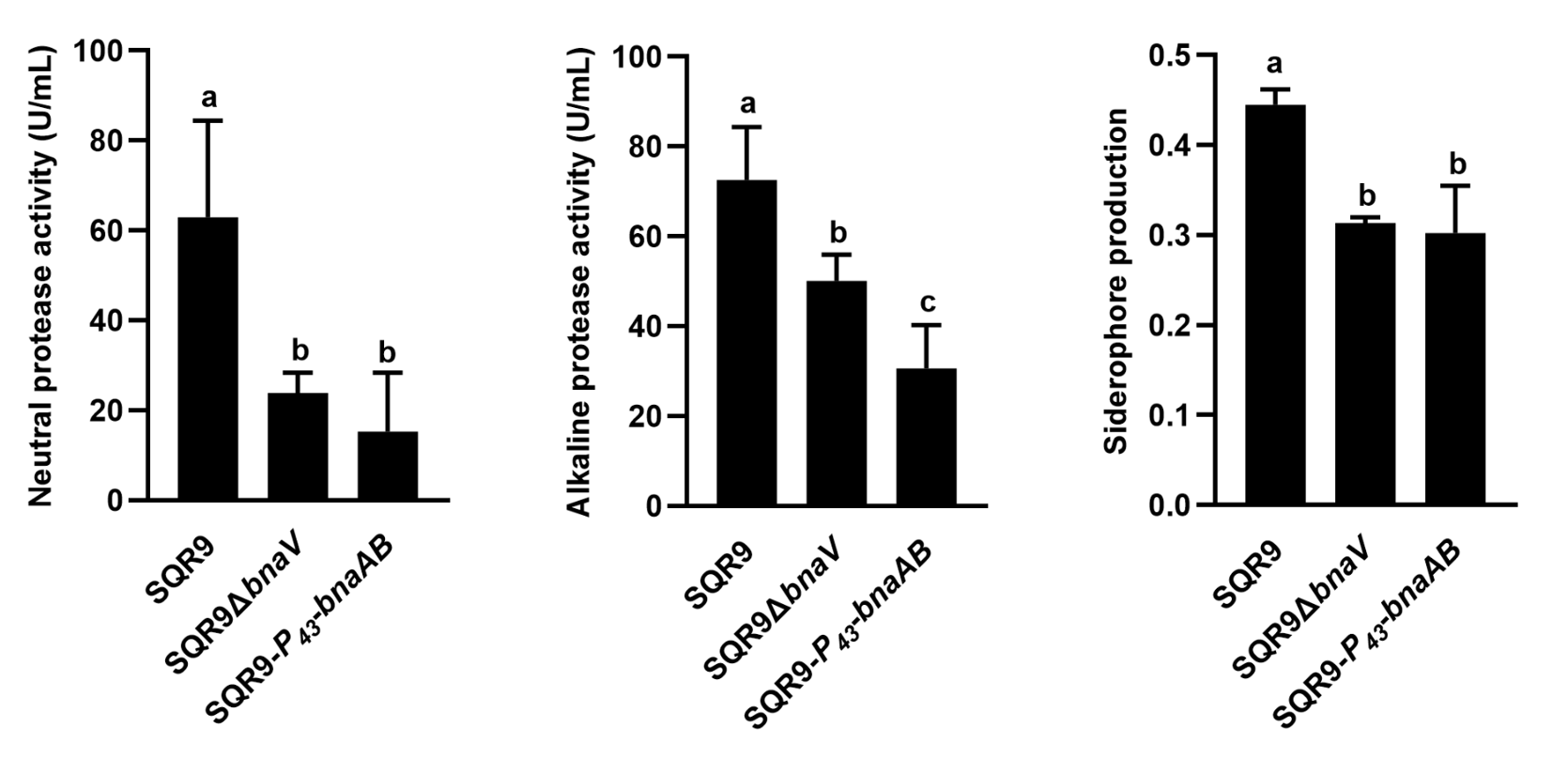

**Fig. S10. Neutral/alkaline protease activity and siderophore production by wild-type SQR9, SQR9Δ*bnaV*, and SQR9-*P_43_-bnaAB*.** Columns with different letters are statistically different according to the Duncan’s multiple range test (*P* < 0.05).

**Table S1. List of strains and plasmids used in this study**

| **Strains or plasmids** | **Description** | **References or sources** |
| --- | --- | --- |
| **Strains** |  |  |
| *Escherichia coli* |  |  |
| *E. coli* Top10 | F-*mcrA*Δ (*mrr*-*hsdRMS*-*mcrBC*) ψ80*lacZ*ΔM15Δ *lacX*74 *nupG* *recA*1 *araD*139Δ (*ara-leu*) 7697 *galE*15 *galK* 16 *rpsL* (Str^R^) end A1λ- | Invitrogen (Shanghai) |
| *E. coli* BL21(DE3) | Ideal for routine T7 expression | Invitrogen (Shanghai) |
| *B. velezensis* | Wild type isolate | (Cao et al., 2011) |
| FZB42 | Wild type isolate | (Chen et al., 2007) |
| SQR9 | Wild type isolate | Lab strain |
| SQR9-*P_eps_*-*gfp* | SQR9 with plasmid pNW33N-*P_eps_*-*gfp*, Cm^r^ | This study |
| SQR9-*P_tasA_*-*gfp* | SQR9 with plasmid pNW33N-*P_tasA_*-*gfp*, Cm^r^ | This study |
| SQR9-*P_fa_*-*gfp* | SQR9 with plasmid pNW33N-*P_fa_*-*gfp*, Cm^r^ | This study |
| SQR9-*P_bnaA_*-*gfp* | SQR9 with plasmid pNW33N-*P_bnaA_*-*gfp*, Cm^r^ | Lab strain |
| SQR9-*P_accDA_*-*gfp* | SQR9 with plasmid pNW33N-*P_accDA_*-*gfp*, Cm^r^ | This study |
| SQR9Δ*spo0A*-*P_eps_*-*gfp* | Δ*spo0A* with plasmid pNW33N-*P_eps_*-*gfp*, Cm^r^, Em^r^ | This study |
| SQR9Δ*spo0A*-*P_tasA_*-*gfp* | Δ*spo0A* with plasmid pNW33N-*P_tasA_*-*gfp*, Cm^r^, Em^r^ | This study |
| SQR9Δ*spo0A*-*P_fa_*-*gfp* | Δ*spo0A* with plasmid pNW33N-*P_fa_*-*gfp*, Cm^r^, Em^r^ | This study |
| SQR9Δ*spo0A*-*P_bnaA_*-*gfp* | Δ*spo0A* with plasmid pNW33N-*P_bnaA_*-*gfp*, Cm^r^, Em^r^ | This study |
| SQR9Δ*spo0A*-*P_accDA_*-*gfp* | Δ*spo0A* with plasmid pNW33N-*P_accDA_*-*gfp*, Cm^r^, Em^r^ | This study |
| SQR9Δ*spo0A*/*spo0A*-*P_eps_*-*gfp* | Δ*spo0A*/*spo0A* with plasmid pNW33N-*P_eps_*-*gfp*, Cm^r^, Em^r^, Spc^r^ | This study |
| SQR9Δ*spo0A*/*spo0A*-*P_tasA_*-*gfp* | Δ*spo0A*/*spo0A* with plasmid pNW33N-*P_tasA_*-*gfp*, Cm^r^, Em^r^, Spc^r^ | This study |
| SQR9Δ*spo0A*/*spo0A*-*P_fa_*-*gfp* | Δ*spo0A*/*spo0A* with plasmid pNW33N-*P_fa_*-*gfp*, Cm^r^, Em^r^, Spc^r^ | This study |
| SQR9Δ*spo0A*/*spo0A*-*P_bnaA_*-*gfp* | Δ*spo0A*/*spo0A* with plasmid pNW33N-*P_bnaA_*-*gfp*, Cm^r^, Em^r^, Spc^r^ | This study |
| SQR9Δ*spo0A*/*spo0A*-*P_accDA_*-*gfp* | Δ*spo0A*/*spo0A* with plasmid pNW33N-*P_accDA_*-*gfp*, Cm^r^, Em^r^, Spc^r^ | This study |
| SQR9-*P_bnaA_*-*gfp*-*P_accDA_*-*rfp* | SQR9 with plasmid pNW33N-*P_bnaA_*-*gfp*-*P_accDA_*-*rfp*, Cm^r^ | This study |
| SQR9-*P_bnaA_*-*gfp*-*P_eps_*-*rfp* | SQR9 with plasmid pNW33N-*P_bnaA_*-*gfp*-*P_eps_*-*rfp*, Cm^r^ | This study |
| SQR9-*P_bnaA_*-*gfp*-*P_fa_*-*rfp* | SQR9 with plasmid pNW33N-*P_bnaA_*-*gfp*-*P_fa_*-*rfp*, Cm^r^ | This study |
| SQR9-*P_bnaA_*-*gfp*-*P_tasA_*-*rfp* | SQR9 with plasmid pNW33N-*P_bnaA_*-*gfp*-*P_tasA_*-*rfp*, Cm^r^ | This study |
| SQR9-*P_fa_*-*gfp*-*P_eps_*-*rfp* | SQR9 with plasmid pNW33N-*P_fa_*-*gfp*-*P_eps_*-*rfp*, Cm^r^ | This study |
| SQR9-*P_fa_*-*gfp*-*P_tasA_*-*rfp* | SQR9 with plasmid pNW33N-*P_fa_*-*gfp*-*P_tasA_*-*rfp*, Cm^r^ | This study |
| SQR9Δ*spo0A* | Em^r^ | Lab strain |
| SQR9Δ*spo0A*/*spo0A* | Em^r^, Spc^r^ | Lab strain |
| SQR9-*P_xyl_*-*accDA* | Replace *P_accDA_* with *P_xyl_*, Spc^r^ | This study |
| SQR9-*P_xyl_*-*accDA-P_fa_-gfp* | SQR9-*P_xyl_*-*accDA* with plasmid pNW33N-*P_fa_*-*gfp*, Cm^r^, Spc^r^ | This study |
| SQR9-*P_xyl_*-*accDA-P_bnaA_-gfp* | SQR9-*P_xyl_*-*accDA* with plasmid pNW33N-*P_bnaA_*-*gfp*, Cm^r^, Spc^r^ | This study |
| SQR9-*P_43_*-*bnaA* | Replace *P_bnaA_* with *P_43_*, Em^r^ | This study |
| SQR9Δ*bnaV* | Em^r^ | Lab strain |
| SQR9Δ*degU* | Zeocin^r^ | Lab strain |
| SQR9Δ*sinI* | Zeocin^r^ | Lab strain |
| SQR9Δ*sinR* | Zeocin^r^ | Lab strain |
| SQR9Δ*abrB* | Zeocin^r^ | Lab strain |
| SQR9Δ*comPA* | Zeocin^r^ | Lab strain |
| **Plasmids** |  |  |
| pNW33n | Cm^r^; *B.subtilis-E.coli* shuttle vector | (Zhou et al., 2017) |
| pNW33N-*P_bnaA_*-*gfp* | pNW33N containing *P_bnaA_*-*gfp* fusion fragment | This study |
| pNW33N-*P_fa_*-*gfp* | pNW33N containing *P_fa_*-*gfp* fusion fragment | This study |
| pNW33N-*P_eps_*-*gfp* | pNW33N containing *P_eps_*-*gfp* fusion fragment | This study |
| pNW33N-*P_tasA_*-*gfp* | pNW33N containing *P_tasA_*-*gfp* fusion fragment | This study |
| pNW33N-*P_accDA_*-*gfp* | pNW33N containing *P_accDA_*-*gfp* fusion fragment | This study |
| pNW33N-*P_bnaA_*-*gfp*-*P_accDA_*-*rfp* | pNW33N containing *P_bnaA_*-*gfp*-*P_accDA_*-*rfp* fusion fragment | This study |
| pNW33N-*P_bnaA_*-*gfp*-*P_eps_*-*rfp* | pNW33N containing *P_bnaA_*-*gfp*-*P_eps_*-*rfp* fusion fragment | This study |
| pNW33N-*P_bnaA_*-*gfp*-*P_fa_*-*rfp* | pNW33N containing *P_bnaA_*-*gfp*-*P_fa_*-*rfp* fusion fragment | This study |
| pNW33N-*P_bnaA_*-*gfp*-*P_tasA_*-*rfp* | pNW33N containing *P_bnaA_*-*gfp*-*P_tasA_*-*rfp* fusion fragment | This study |
| pNW33N-*P_fa_*-*gfp*-*P_eps_*-*rfp* | pNW33N containing *P_fa_*-*gfp*-*P_eps_*-*rfp* fusion fragment | This study |
| pNW33N-*P_fa_*-*gfp*-*P_tasA_*-*rfp* | pNW33N containing *P_fa_*-*gfp*-*P_tasA_*-*rfp* fusion fragment | This study |
| pMAL-c5X-*spo0A* | pMAL-c5X containing sequence encoding Spo0A, Amp^r^ | This study |

**Table S2. List of primers used in this study.**

| **Name** | **Sequence (5’-3’)** |
| --- | --- |
| pN-*P_tasA_-gfp*-1F | CGAGCTCGTATTGCTTGACTGCTTCG |
| pN-*P_tasA_-gfp*-1R | AGTTCTTCTCCTTTACTCATCTTTTTTTCCATTCGCAAC |
| pN-*P_tasA_-gfp*-2F | GTTGCGAATGGAAAAAAAGATGAGTAAAGGAGAAGAACT |
| pN-*P_tasA_-gfp*-2R | CCGCTCGAGTTATTTGTATAGTTCATCCAT |
| pN-*P_eps_-gfp*-1F | CGAGCTCATTGGATGGACGGATTTG |
| pN-*P_eps_-gfp*-1R | AGTTCTTCTCCTTTACTCATTAATTCTTTAAAACTCATATTC |
| pN-*P_eps_-gfp*-2F | GAATATGAGTTTTAAAGAATTAATGAGTAAAGGAGAAGAACT |
| pN-*P_fa_-gfp*-1F | CGAGCTCAGAGATGCATGATTTAACAT |
| pN-*P_fa_-gfp*-1R | AGTTCTTCTCCTTTACTCAT GAACACTTCAAAAATTTCTT |
| pN-*P_fa_-gfp*-2F | AAGAAATTTTTGAAGTGTTCATGAGTAAAGGAGAAGAACT |
| pN-*P_accDA_-gfp*-1F | CGAGCTCACAAGACAGCTCACGGTTAT |
| pN-*P_accDA_-gfp*-1R | AGTTCTTCTCCTTTACTCATATGATTACCTCCCTTTTGTG |
| pN-*P_accDA_-gfp*-2F | CACAAAAGGGAGGTAATCATATGAGTAAAGGAGAAGAACT |
| pN-*P_accDA_*-*rfp*-1F | CGAGCTC ACAAGACAGCTCACGGTTAT |
| pN-*P_accDA_*-*rfp*-1R | CTCGCCCTTGCTCACCATATGATTACCTCCCTTTTGTG |
| pN-*P_accDA_*-*rfp*-2F | CACAAAAGGGAGGTAATCATATGGTGAGCAAGGGCGAG |
| pN-*P_accDA_*-*rfp*-2R | CCGCTCGAGCTACTTGTACAGCTCGTCCA |
| pN-*P_bnaA_*-*gfp*-*P_eps_*-*rfp*-1F | CGAGCTC TGAAGAATATGTTGAAACAGTT |
| pN-*P_bnaA_*-*gfp*-*P_eps_*-*rfp*-1R | TGGACGAGCTGTACAAGTAGTTATTTGTATAGTTCATCCATGC |
| pN-*P_bnaA_*-*gfp*-*P_eps_*-*rfp*-2F | GCATGGATGAACTATACAAATAACTACTTGTACAGCTCGTCCA |
| pN-*P_bnaA_*-*gfp*-*P_eps_*-*rfp*-2R | CCGCTCGAG ATTGGATGGACGGATTTG |
| pN-*P_fa_*-*rfp*-1F | CGAGCTCAGAGATGCATGATTTAACAT |
| pN-*P_fa_*-*rfp*-1R | CTCGCCCTTGCTCACCATGAACACTTCAAAAATTTCTT |
| pN-*P_fa_*-*rfp*-2F | AAGAAATTTTTGAAGTGTTCATGGTGAGCAAGGGCGAG |
| pN-*P_fa_*-*rfp*-2R | CCGCTCGAGCTACTTGTACAGCTCGTCCA |
| pN-*P_bnaA_*-*gfp*-*P_tasA_*-*rfp*-1F | CGAGCTCTGAAGAATATGTTGAAACAGTT |
| pN-*P_bnaA_*-*gfp*-*P_tasA_*-*rfp*-1R | TGGACGAGCTGTACAAGTAGTTATTTGTATAGTTCATCCATGC |
| pN-*P_bnaA_*-*gfp*-*P_tasA_*-*rfp*-2F | GCATGGATGAACTATACAAATAACTACTTGTACAGCTCGTCCA |
| pN-*P_bnaA_*-*gfp*-*P_tasA_*-*rfp*-2R | CCGCTCGAGGTATTGCTTGACTGCTTCG |
| pN-*P_fa_*-*gfp*-*P_eps_*-*rfp*-1F | CGAGCTC AGAGATGCATGATTTAACATT |
| pN-*P_fa_*-*gfp*-*P_eps_*-*rfp*-1R | TGGACGAGCTGTACAAGTAGTTATTTGTATAGTTCATCCATGC |
| pN-*P_fa_*-*gfp*-*P_eps_*-*rfp*-2F | GCATGGATGAACTATACAAATAACTACTTGTACAGCTCGTCCA |
| pN-*P_fa_*-*gfp*-*P_eps_*-*rfp*-2R | CCGCTCGAG ATTGGATGGACGGATTTG |
| pN-*P_fa_*-*gfp*-*P_tasA_*-*rfp*-1F | CGAGCTC AGAGATGCATGATTTAACATT |
| pN-*P_fa_*-*gfp*-*P_tasA_*-*rfp*-1R | TGGACGAGCTGTACAAGTAGTTATTTGTATAGTTCATCCATGC |
| pN-*P_fa_*-*gfp*-*P_tasA_*-*rfp*-2F | GCATGGATGAACTATACAAATAACTACTTGTACAGCTCGTCCA |
| pN-*P_fa_*-*gfp*-*P_tasA_*-*rfp*-2R | CCGCTCGAG GTATTGCTTGACTGCTTCG |
| *P_xyl_*-*accDA*-UF | GAGTAGCTGAGCCTTGTAAAG |
| *P_xyl_*-*accDA*-UR | CGTTACGTTATTAGTTATCAGAACATTGTTAACCTGGT |
| *P_xyl_*-*accDA*-SpcF | ACCAGGTTAACAATGTTCTGATAACTAATAACGTAACG |
| *P_xyl_*-*accDA*-SpcR | TAAGTGTTACCCCTATAAGTTAGGTTACGTATAATGTATGCTATA |
| *P_xyl_*-*accDA*-*P_xyl_*F | TATAGCATACATTATACGTAACCTAACTTATAGGGGTAACACTTA |
| *P_xyl_*-*accDA*-*P_xyl_*R | TGTGAATATATCCTTTAACAAGTTTTCTGACTCATATCGAACATT |
| *P_xyl_*-*accDA*-BF | AATGTTCGATATGAGTCAGAAAACTTGTTAAAGGATATATTCACA |
| *P_xyl_*-*accDA*-BR | CCGTGCTTTAATAAAAAT |
| *P_43_*-*bnaA*-UF | GCTAAGAATGGTAAAGCGTA |
| *P_43_*-*bnaA*-UR | ACTTTTCGGGGAAATGTGTTAAACCTTATATAGTCGATAACC |
| *P_43_*-*bnaA*-EmF | GGTTATCGACTATATAAGGTTTAACACATTTCCCCGAAAAGT |
| *P_43_*-*bnaA*-EmR | CGAAAACATACCACCTATCATTATTTCCTCCCGTTAAATA |
| *P_43_*-*bnaA*-*P_43_*-F | TATTTAACGGGAGGAAATAATGATAGGTGGTATGTTTTCG |
| *P_43_*-*bnaA*-*P_43_*-R | TTCAAAATGAATTTTTCCATGTGTACATTCCTCTCTTACCTA |
| *P_43_*-*bnaA*-BF | TAGGTAAGAGAGGAATGTACACATGGAAAAATTCATTTTGAA |
| *P_43_*-*bnaA*-BR | TCCTTTATTAAGAAGAGCTTG |
| pMAL-spo0A-F | ACGCGTCGACGTGGAGAAAATTAAAGTTTGT |
| pMAL-spo0A-R | CGCGGATCCTTAGTGGTGGTGGTGGTGGTGCGAAGCTTTATGCTCCA |

**Movie S1. Dynamic observation of extracellular polysaccharides (EPS)-producing cells and dead cells in *B. velezensis* SQR9 community during biofilm formation.** Colony cells of SQR9-*P_eps_*-*gfp* was stained with propidium iodide (PI, a red-fluorescent dye for labeling dead cell) for 15 min, and then visualized by a CLSM to monitor the distribution of fluorescence signal from reporter and the PI dye. The movie is consisted of pictures obtained at 0, 10, and 20 min after treatment (each picture retains 2 secs). The bar represents 5 μm.

**Movie S2. Dynamic observation of TasA fibers-producing cells and dead cells in *B. velezensis* SQR9 community during biofilm formation.** Colony cells of SQR9-*P_tapA_*-*gfp* was stained with PI for 15 min, and then visualized by a CLSM to monitor the distribution of fluorescence signal from reporter and the PI dye. The movie is consisted of pictures obtained at 0, 5, 10, 15, and 20 min after treatment (each picture retains 2 secs). The bar represents 5 μm.

**Movie S3. Dynamic observation of bacillunoic acids (BAs)-producing cells and dead cells in *B. velezensis* SQR9 community during biofilm formation.** Colony cells of SQR9-*P_bnaF_*-*gfp* was stained with PI for 15 min, and then visualized by a CLSM to monitor the distribution of fluorescence signal from reporter and the PI dye. The movie is consisted of pictures obtained at 0, 5, 10, 15, and 20 min after treatment (each picture retains 2 secs). The bar represents 5 μm.

**Movie S4. Dynamic observation of bacillunoic acids (BAs)-immunity cells and dead cells in *B. velezensis* SQR9 community during biofilm formation.** Colony cells of SQR9-*P_bnaAB_*-*gfp* was stained with PI for 15 min, and then visualized by a CLSM to monitor the distribution of fluorescence signal from reporter and the PI dye. The movie is consisted of pictures obtained at 0, 5, 10, 15, and 20 min after treatment (each picture retains 2 secs). The bar represents 5 μm.
